# The Persistence Potential of Mobile Genetic Elements

**DOI:** 10.1101/2020.03.03.975128

**Authors:** Teng Wang, Lingchong You

## Abstract

Mobile genetic elements (MGEs), such as plasmids, phages, and transposons, play a critical role in mediating the transfer and maintenance of diverse traits and functions in microbial communities. This role depends on the ability of MGEs to persist. For a community consisting of multiple populations transferring multiple MGEs, however, the conditions underlying the persistence of these MGEs are poorly understood. Computationally, this difficulty arises from the combinatorial explosion associated with describing the gene flow in a complex community using the conventional modeling framework. Here, we describe an MGE-centric framework that makes it computationally feasible to analyze such transfer dynamics. Using this framework, we derive the persistence potential: a general, heuristic metric that predicts the persistence and abundance of any MGEs. We validate the metric with engineered microbial consortia transferring mobilizable plasmids and quantitative data available in the literature. Our modeling framework and the resulting metric have implications for developing a quantitative understanding of natural microbial communities and guiding the engineering of microbial consortia.

## Introduction

Mobile genetic elements (MGEs), including plasmids, transposons and phages are major components of the metagenome of microbes^1,2^. Hundreds of plasmids have been identified in diverse microbial communities from rat cecum^3^, cow rumen^4^, sludge^5^, and marine water^6^. In the reference genomes of the human gut microbiome, ~16,000 genes are identified as mobile^7^. MGEs encode diverse biological functions including metabolic capabilities^8^, pathogenic virulence^9^, or traits to cope with environmental stresses^10–12^. The interplay between MGEs and the core genomes of the host cells shapes the evolution of microbial communities^13^.

The ability for an MGE to persist in a microbial community can influence the dynamics, function, and even survival of the community^14,15^. Promotion or suppression of MGE persistence, depending on the context, has applications in medicine, biosafety, and biotechnology. For instance, eliminating an MGE encoding resistance to an antibiotic will sensitize a community to the antibiotic^16^, and this resistance reversal can enable more effective use of antibiotics^17^. There is always a biosafety concern about the risk associated with the spread of synthetic genetic constructs into the environment^18,19^. To reduce the risk, the use of genetic vectors with restricted capability to be maintained outside of the laboratory has been proposed^18,19^. In biotechnology, plasmid instability is a major impediment to the large-scale production of recombinant protein products^20,21^. The expression of recombinant proteins encoded by the plasmids are usually burdensome to the cell metabolism, and, as a consequence, the plasmid-carrying cells can be outcompeted by the faster-growing plasmid-free populations. A strategy to promote plasmid persistence could overcome this limitation.

The quantitative studies of MGE persistence and abundance in microbial communities is challenging due to the lack of an effective theoretical framework^22,23^. Since the 1970s, population-biology models have been developed to predict the persistence of a single MGE in a single species^16,24–28^. However, microbes in nature often live in complex communities consisting of diverse species and mobile elements^7,29,30^. The limited scope of past quantitative analyses is due in part to the challenge associated with modeling complex communities using the conventional modeling framework, which we refer to as ‘subpopulation-centric framework’ (SCF). In SCF, a population carrying a particular combination of MGEs is considered a unique subpopulation that requires one ordinary differential equation (ODE) to describe. Modelling a community containing two species and two mobile elements requires eight ODEs, each describing one subpopulation. The model complexity increases combinatorially with the number of MGEs (Fig. 1A and B). For example, the marine microbiome in a bottle of sea water is estimated to contain ~160 species^30^ and ~180 plasmids^6^. In SCF, ~2.4 × 10^56^ ODEs and ~2.7 × 10^114^ parameters are needed to model the gene flow dynamics of this community. Therefore, it is currently impossible to model a system in which many MGEs are involved.

**Fig. 1.**
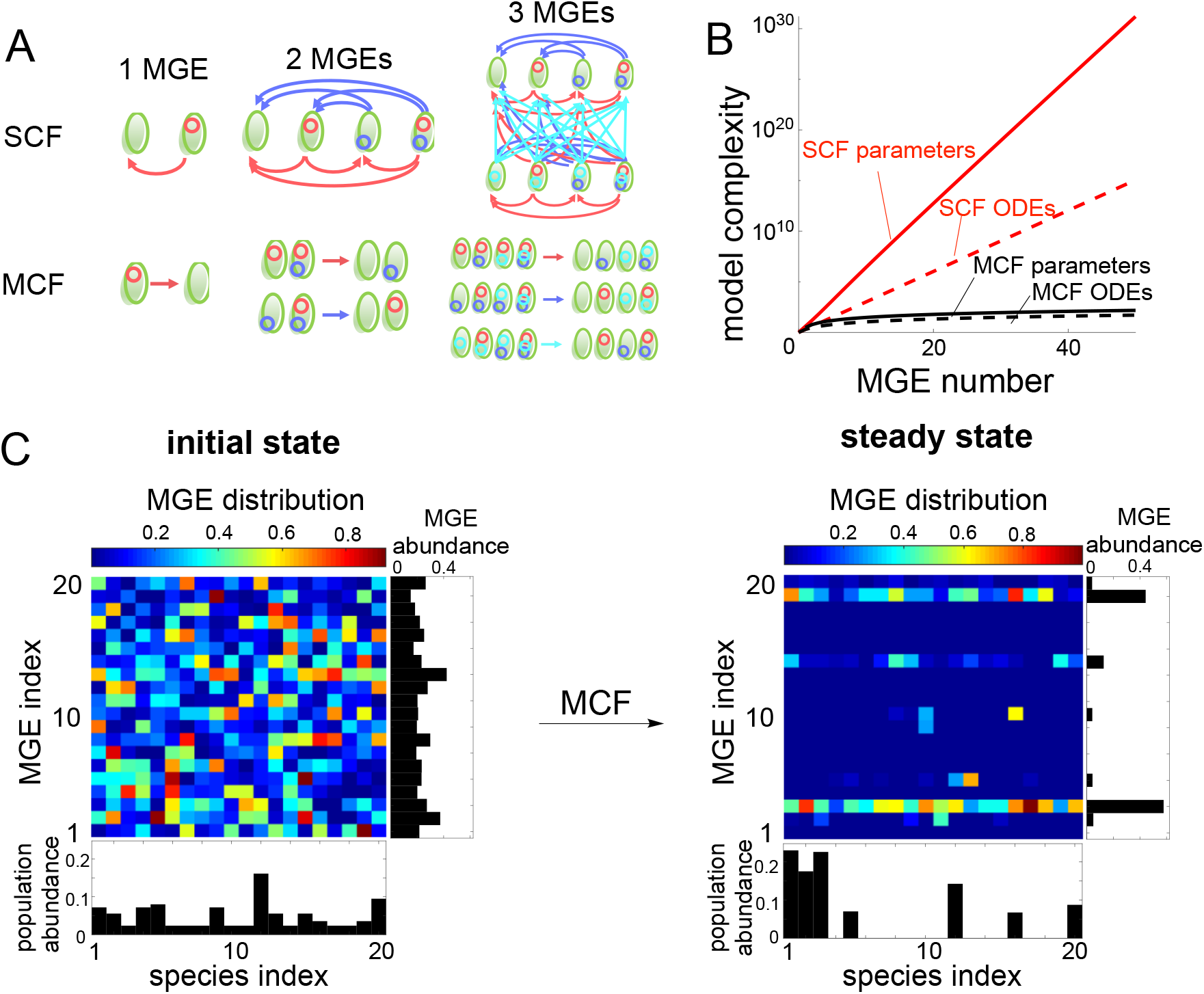
An MGE-centric framework (MCF) to model horizontal gene transfer in microbial communities. **(A) Comparison between the subpopulation-centric framework (SCF) and MCF for one species transferring 1-3 mobile genetic elements (MGEs).** The arrows represent the MGE transfer from the donor to the recipient. **(B) Model complexity of single-species communities as a function of the number of MGEs.** The model complexity refers to the number of ordinary differential equations (ODEs) or the number of parameters required in each model. **(C) Simulation of the dynamics of a community consisting of 20 species transferring 20 MGEs.** Different species or MGEs are distinguished by the indexes. A random set of parameter values were used. The initial distributions (left) of population sizes and MGE abundances were also random. The right panel shows the steady-state distributions of the MGEs across different species, quantified as the relative abundance of MGE in each species. The relative abundance of MGE *j* in species *i* is calculated as the fraction of species *i* cells that contains MGE *j* relative to the total number of species *i* cells. The total abundances of each species and each MGE in the entire community are shown as bars.

## Results

### An MGE-centric framework to model gene flow

To overcome the challenge associated with SCF, we developed an MGE-centric framework (MCF) that focuses on the overall abundance of each MGE in the community by accounting for the average fitness effect of MGEs to the host. To illustrate the key concepts, consider the transfer of *n* types of mobilizable plasmids in a community consisting of *m* species. Let *s*_*i*_ (*i* = 1,2, …, *m*) be the abundance of *i*-th species and *p*_*ij*_ (*j* = 1,2, …, *n*) the abundance of the *i*-th species carrying the *j*-th plasmid (Fig. S1A). The dynamics can be approximately described by two groups of ODEs (see *Supplementary Information* section 2.1 for detailed derivation and justification)

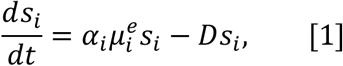

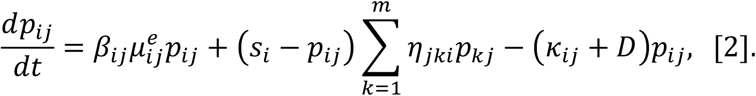

Eq.1 describes the collective growth and dilution of *s*_*i*_, where 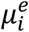 is the effective growth rate of the species, *D* is the dilution rate, and *α*_*i*_ represents the combined growth effect of all the plasmids carried by the species. In Eq.2, the first term describes the growth of *p*_*ij*_. 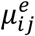 is the effective growth rate; it differs from 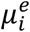 due to the growth effect of the *j*-th plasmid. *β*_*ij*_ is the combined growth effect of the other plasmids. In the second term, *η*_*iki*_ is the conjugation efficiency when the plasmid is transferred from the *k*-th species to the *i*-th species. *s*_*i*_ − *p*_*ij*_ is the total abundance of subpopulations of *s*_*i*_ not carrying the *j*-th plasmid. The third term describes the plasmid loss due to segregation error (at a rate constant of *κ*_*ij*_) and dilution.

In this formulation, *p*_*ij*_/*s*_*i*_ indicates the relative abundance of the *j*-th plasmid in the *i*-th species; 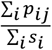 is the relative abundance of the *j*-th plasmid in the entire community. Because of the formulation of *α*_*i*_, *β*_*ij*_, 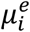 and 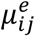, this framework is applicable for describing arbitrary microbial communities transferring multiple plasmids. In particular, the interactions between populations are accounted for by the appropriate formulation of 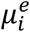 and 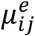 plasmid incompatibility can be accounted for by adapting the formulations of *β*_*ij*_ and the conjugation terms (Fig. S1C, see *Supplementary Information* section 2.1.3).

For a community consisting of *m* species transferring *n* plasmids, the SCF requires *m* ∙ 2^M^ ODEs and approximately *nm*^2^ ∙ 2^2*n*-2^ parameters, whereas our MCF only needs *m*(*n* + 1) ODEs and approximately *nm*^N^ parameters (Fig. 1A and B, Table S2, see *Supplementary Information* section 2.2). This simplification renders our framework capable of predicting the dynamics of species composition as well as the distribution patterns of each plasmid (Fig. 1C). In particular, we modeled a community consisting of 200 species transferring 200 plasmids. The conventional SCF requires about 3.2 × 10^62^ ODEs and 5.2 × 10^126^ parameters, which is impractical to construct. In contrast, this community can be modeled by 4.0 × 10^4^ ODEs and 8.1 × 10^6^ parameters using MCF, which is feasible to both construct and calculate (Fig. S1B). Although acquiring all the parameters in such complex communities remains challenging due to the technical limitations ^31^, MCF enables the theoretical analysis of the dynamics and persistence conditions of MGEs.

This drastic reduction in model complexity is made possible by lumping multiple distinct subpopulations into an average one and then accounting for the average fitness effect of each plasmid. In particular, *s*_*i*_ lumps all subpopulations carrying different combinations of plasmids or no plasmids and *p*_*ij*_ lumps all subpopulations carrying the *j*-th plasmid. If the plasmids do not have any growth effects, the lumping is exact; the MCF is equivalent to SCF. In general, however, plasmids can confer burden or benefit, which will cause deviation between the results computed from these two frameworks. To evaluate this discrepancy, we conducted numerical simulations on communities transferring one and two plasmids. Testing higher number of plasmids in SCF is computationally prohibitive due to the combinatorial explosion. Indeed, simulation results suggest that the fitness effects of the plasmids are the main factors that determine the discrepancy. Within a reasonably wide range of fitness costs, the predictions of these two models match well, and the smaller the fitness effects, the smaller the discrepancy. (Fig. S2, see *Supplementary Information* section 2.3). In general, we note that both frameworks represent approximations of real biological systems. The ultimate test of each framework should be from experiments.

The same framework is applicable to the analysis of other MGEs, including phages and transposons. Temperate phages exit the host cells by budding or secretion ^32^, without killing the host. This transfer process can be described by MCF by interpreting conjugation efficiencies (*η*’s) as phage transfer rates. Lytic phages, which cause host death and lysis during release ^27,33,34^, can be described by the MCF with suitable modifications (see *Supplementary Information* section 2.4). Transposons are genetic elements that locate on chromosomes, plasmids or phages ^1^, and their intercellular transfer of transposons is mediated by plasmid conjugation or phage transduction ^1,35^. The MCF is directly applicable to transposons (see *Supplementary Information* section 2.4).

### The persistence potential of an MGE

Our framework makes it feasible to develop a criterion of MGE persistence for complex communities. We first considered an idealized community, where each species has the same set of kinetic parameters and equal abundance (Fig. 2A). Because of its symmetry, this community allows the analytical derivation of a metric (*ω*) that determines the persistence and steady-state abundance of the plasmids relative to the total cell density (Fig. 2A, also see *Supplementary Information* section 2.5)

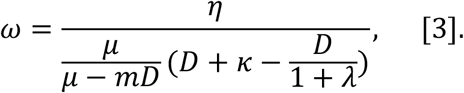

**Fig. 2.**
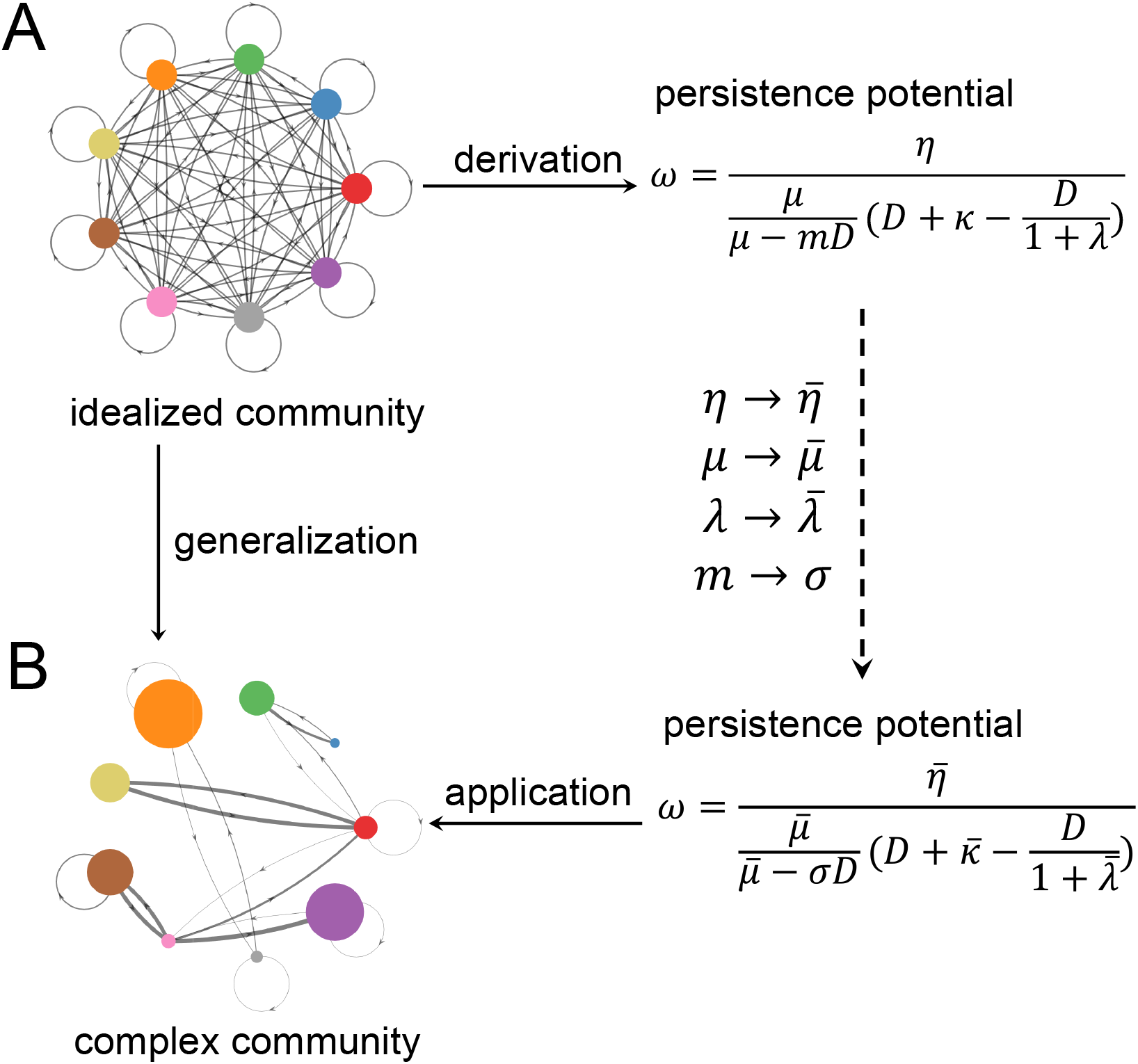
Developing the persistence potential (*ω*) of mobile genetic elements (MGEs). **(A) Derivation of *ω* for an idealized microbial community**. Here, the community is fully symmetric in terms of parameters and population sizes. Each colored circle represents a constituent species; the area of the circle is proportional to the species size. Each directed arrow represents the transfer of an MGE, with the arrow thickness indicating the transfer rate. In this case, *ω* can be analytically derived. **(B) Generalization of** *ω*. We generalized the formulation of *ω* by replacing each parameter with its weighted average, which factors in the heterogeneity of species size. The weighted averages of *μ*, *κ*, and *λ* were calculated by 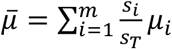, 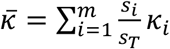, 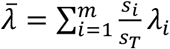, where *m* is the total number of species and *s*_*i*_ represents the abundance of *i*-th species. *s*_Z_ is the total abundance of all the populations: 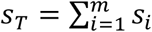. Both donor and recipient cell densities contribute to the conjugation efficiency; thus, the weighted average of *η* was calculated by 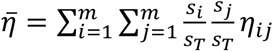, where *s*_*j*_ represents the abundance of the donor species, *s*_4_ the recipient species, and *η*_*ij*_ the transfer rate from the donor to the recipient.

We term this metric the **persistence potential** for the MGE. In this idealized community, a burdensome MGE (*λ* >0) persists if and only if *ω* > 1. If the mobile element is beneficial (*λ* <0), which leads to *ω* <0, the element will also persist.

In general, the constituent species in a community are not symmetric (Fig. 2B). Their abundances are different from each other; they have different growth rates; and they transfer plasmids at different rates. For such communities, deriving an analytical solution is not possible. We thus took a heuristic approach to generalize the idealized metric. We kept its derived formulation but replaced each parameter with the weighted average of the corresponding parameter in the general community accounting for the relative abundances of different populations: 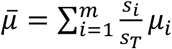, 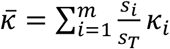, 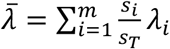 and 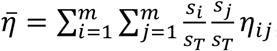, where *s*_*T*_ is the total abundance of all the populations: 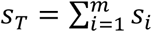 (Fig. 2). Correspondingly, we replace the number of species, *m*, with the community diversity *σ*, which is calculated through 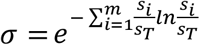. The general form of persistence potential becomes:

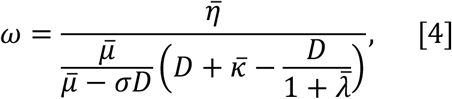

where 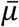, 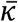, 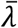, 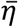 and *D* stand for the weighted averages of species growth rate, plasmid loss rate, fitness cost, horizontal transfer rate, and dilution rate, respectively (Fig. 3A and B).

To evaluate the predictive power of the generalized persistence potential, we performed numerical simulations with 2,000 sets of randomized parameters in communities consisting of 5-100 species and 1-50 plasmids. Our randomization process produced communities with various settings that mimic the diversity of the natural genetic-exchange communities. To ensure there were adequate numbers of species coexisting at steady states, the community was divided into multiple coexisting niches. Each niche had its own carrying capacity, and different species within the same niche competed with each other (Fig. S1D). The dynamics of each community was simulated for up to ~30,000 hours until it became unchanged, and we treated it as ‘steady state’. The fractions of the MGE-carrying cells were then calculated with respect to *ω* values. Our simulation results suggest that the general metric, despite its heuristic nature, remains a robust predictor on whether and to what extent an MGE can persist, with a transition at *ω* = 1 (Fig. 3C). When 0 < *ω* <1, the abundance of the MGE is close to 0; when *ω* > 1, the abundance of the MGE increases monotonically with *ω*. The data points computed from the randomized parameter sets approximately collapsed into a single curve, underscoring the general predictive power of *ω*.

**Fig. 3.**
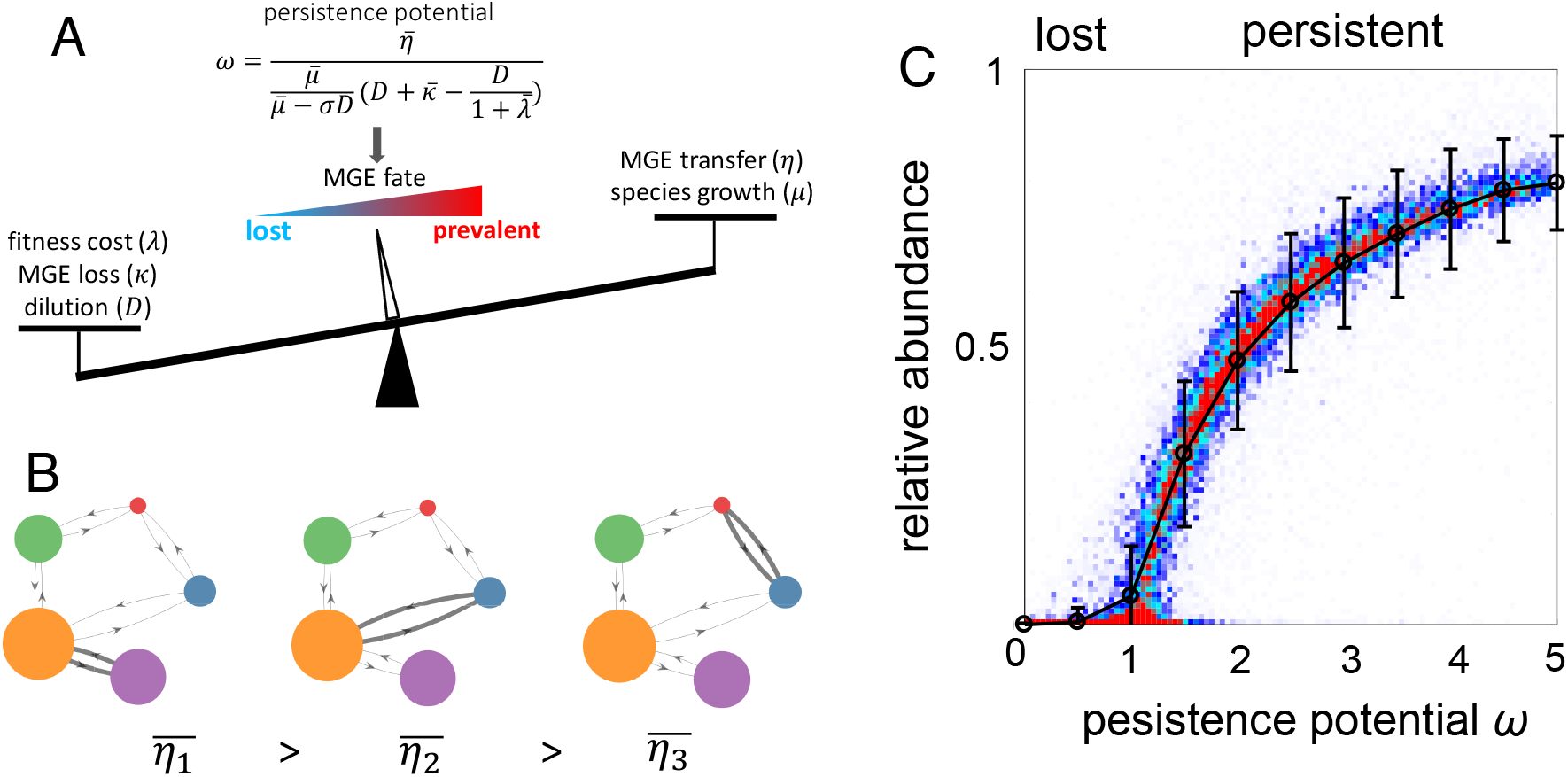
Persistence potential ω predicts the maintenance and abundance of mobile genetic elements (MGEs). **(A) ω accounts for the contribution from multiple parameters.** The MGE horizontal transfer and population growth promote the *ω* value, while fitness cost, MGE loss, and dilution suppress the *ω* value. **(B) The weighted average transfer rates 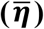 are calculated with respect to population fractions**. A community of five populations and one mobile element is illustrated as an example. The arrows represent the horizontal transfer of the MGE, and the width of the arrow represents the magnitude of the transfer rate. The filled circles represent the populations, with their areas corresponding to the abundance. With equivalent conjugation rates, the conjugation pairs with higher donor or recipient abundance contribute more to 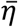. **(C) ω is a robust predictor of the steady-state abundance of MGEs**. 2,000 simulations were performed with 5-100 populations, 1-50 MGEs, and randomized parameters in the range of 0.4 ≤ *μ* ≤ 0.8,0.001 ≤ *D* ≤ 0.005, 0 ≤ *κ* ≤ 0.002, 0 ≤ *λ* ≤ 0.2. For each simulation, the communities were assembled into a random number of niches. Within each niche, populations compete with each other. Each simulation was initialized with random abundances of populations and MGEs. The steady-state persistence potential *ω* of each MGE and its relative abundance in the entire community were then calculated. The *ω* range is divided into multiple bins with widths of 0.5. The standard deviations of the relative abundance in each bin are shown as error bars.

In addition to assuming symmetry, the basic form of *ω* (Eq. 4) is derived by assuming that the system has reached steady state. However, this assumption is not required for the approximate predictive power of *ω*. In particular, in the numerical simulation described above, the *ω* values have similar predictive power for the MGE abundance well before the system has reached steady state (Fig. S3). This result underscores the general predictive power of *ω* and its applicability to experimental systems, which may not be at steady state.

### Experimental validation of the persistence potential

To test the predictive power of the persistence potential, we engineered eight communities transferring mobilizable plasmids. Community 1 through 7 were constructed from three *E. coli* strains (denoted X, B, and R). Strain B expresses BFP on the chromosome, R expresses dTomato, and strain X is not fluorescent. The mobilizable plasmid K was transferred among the strains and expresses GFP constitutively. Using flow cytometry, this system allows the simultaneous quantification of plasmid abundance (with GFP) and population compositions (with BFP and dTomato) (Fig. S4A). Community 8 contained two *E. coli* strains (MG1655 and DH5α) and five conjugative plasmids (F’, PCU1, R388, R6K, and RP4). The community composition and plasmid abundance were quantified by selective plating.

After plasmids were introduced into the communities, dilutions of four different ratios (10^3^ ×, 10^4^ ×, 10^5^ ×, 10^6^ ×) were performed every 24 hours to maintain the growth. We monitored the population dynamics daily over the following 15 days and obtained the fractions of each population and the plasmid (Fig. S4B-H, Fig. S5C). We measured the conjugation rates and fitness costs (Fig. S4I and J, Fig. S5A and B) and also estimated the average dilution rate *D* and plasmid segregation loss rate. With these parameters, we were able to calculate the plasmid persistence potential in each community. The results were well matched to the predicted pattern, suggesting that *ω* values determine plasmid abundance in microbial communities (Fig. 4A).

**Fig. 4.**
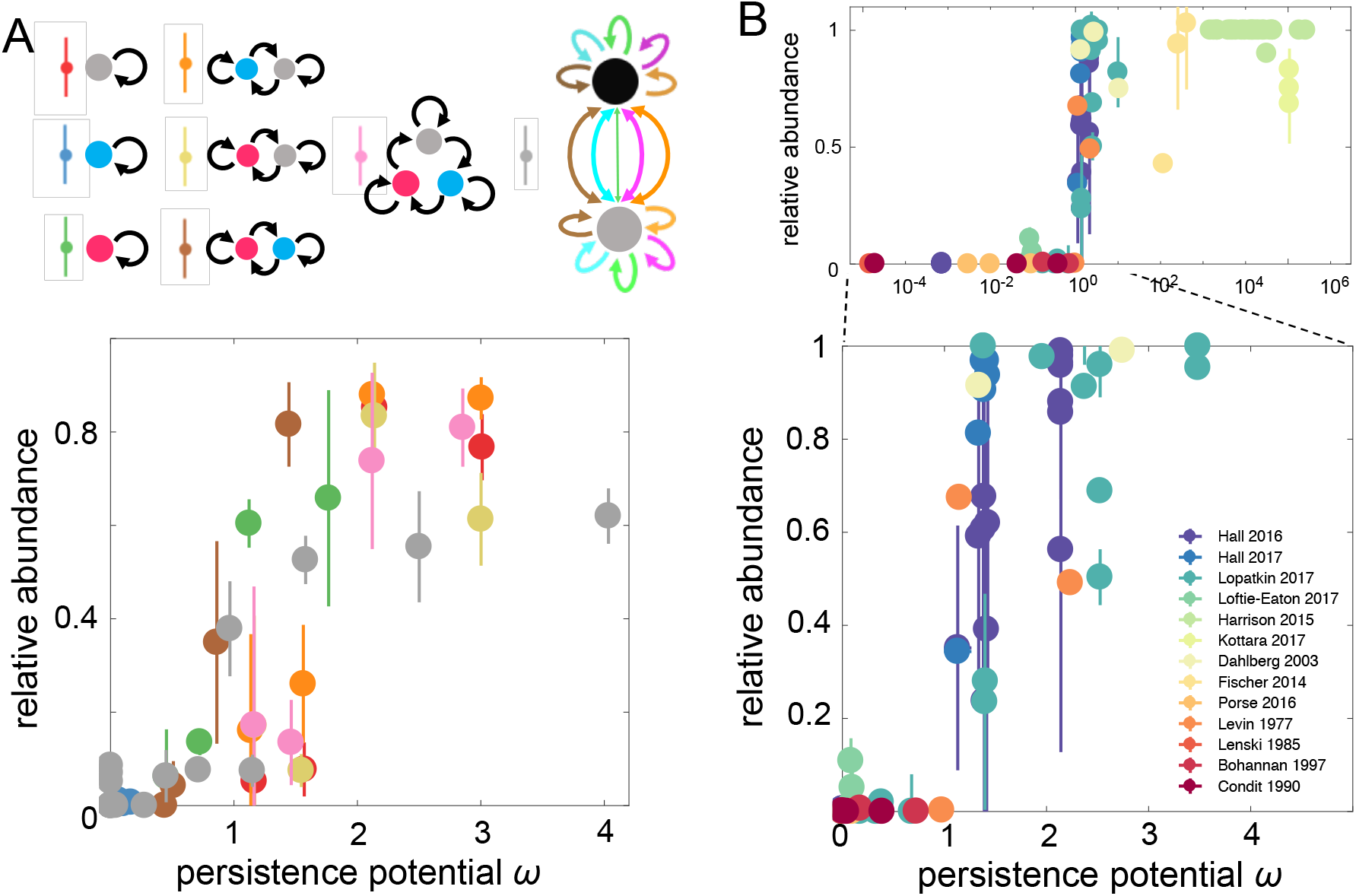
Experimental validation of the predictive power of ω. **(A) Experimental test with eight synthetic communities.** The first seven communities (upper panel) transferring a single MGE were constructed using three engineered *E. coli* strains. Strain B (blue) expresses BFP chromosomally; strain R (red) expresses dTomato; and strain Y (gray) does not express a fluorescent marker. We assembled three single-population communities, two pairwise communities, and one three-population community. Black arrows indicate the plasmid transfer. Dilutions of four different ratios (10^3^ ×, 10^4^ ×, 10^5^ ×, 10^6^ ×) were performed every 24 hours to maintain the growth. The relative abundances and persistence potentials of plasmid K in the seven communities at the end of the experiments (day 15) are shown in the lower panel. The error bars represent the standard deviations of three replicates. The eighth community transferring multiple plasmids was composed of two *E. coli* strains, MG1655 and DH5α. Five self-mobilizable plasmids, F’ (lncF, Tet^R^), PCU1 (lncN, Amp^R^),R388 (lncW, Tm^R^), R6K (lncX, Strp^R^), and RP4 (lncP, Kan^R^), were transferring within the community. The composition of the community was determined by blue-white screening on X-gal plates. The relative abundance of each plasmid was determined by selective plating with the corresponding antibiotics. Four different ratios (10^3^ ×, 10^4^ ×, 10^5^ ×, 10^6^ ×) were performed every 24 hours. The relative abundances and persistence potentials of the five plasmids at the end of the experiments (day 15) are shown in the lower panel. The error bars represent the standard deviations of three replicates. **(B) Evaluation of literature data.** Data were extracted and reanalyzed from 13 previous studies (Tables S3-S19). The error bars represent the standard deviations of multiple replicates. The distribution is shown in the logarithmic scale (upper panel) or linear scale of *ω* (lower panel).

To examine the general applicability of the metric, we reanalyzed the data from 13 previous studies that have provided sufficient measurements on the kinetic parameters and abundance of plasmids^16,36–43^, transposons^35^ and bacteriophages^33,34,44^. The microbial communities analyzed in these studies covered one, two, or three populations, and up to three MGEs. We collected a total of 92 data points (see *Supplementary Information* section 3) covering a *ω* range from 1.46 × 10^−5^ to 2.62 × 10^5^ (Fig. 4B). These data confirm the predictive power of *ω*: in general, the MGE persists when *ω* > 1, and its relative abundance increases with *ω*. In contrast, when *ω* < 1, the MGE tend not to persist, with its relative abundance close to 0.

## Discussion

Our work addresses two fundamental challenges facing the quantitative analysis of gene flow dynamics in microbial communities. First, it has been impractical to simulate a complex community in which more than dozen MGEs are transferred. This challenge is resolved by using our MGE-central modeling framework through drastic dimension reduction in model formulation. Second, the MCF enables the heuristic derivation of a metric that predicts the persistence and abundance of *any* MGEs in a microbial community based on its kinetic parameters and the community composition. Both the modeling framework and the derived MGE persistence potential have implications for future efforts to understand, control, and exploit horizontal gene transfer dynamics in microbial communities.

For instance, the persistence potential *ω* predicts that the abundance of an MGE in a microbial community is sensitive to the average growth rate of the constituent populations, especially when the growth rate is close to the system dilution rate. Consistent with this notion, nutrient enrichments, which in general increase community average growth rate, have been shown to increase the relative abundance of plasmids^45^ and bacteriophages^44^. In the inflamed mouse gut, the growth of commensal Enterobacteriaceae such as *E.coli* is boosted by the nitrate generated as a by-product of the host inflammatory response^46^; this transient boom of Enterobacteriaceae was shown to correlate with an increase in transconjugant abundance^47,48^. In both situations, an increase in the growth rate can promote persistence by increasing the persistence potential directly (through a decrease of its denominator term, Eq2) or indirectly (through increasing the conjugation efficiency)^49^. Conversely, our metric suggests that modulating the overall growth rate of gut microbiota by nutrient supply or its dilution by water inflow could be effective strategies to regulate the abundance of the mobile gene pool in the human gut^50,51^.

Our metric also predicts the influence of bacterial community composition on the MGE persistence potential. For example, the presence of bacterial species with strong efficiencies for interspecific MGE transfers is predicted to promote the average persistence potential of the MGEs. Indeed, for bacteria with poor conjugation efficiencies, coculturing them with efficient donors has been shown to enhance plasmid transfer and maintenance^36,52^. The human gut is equipped with an enormous diversity of microbes^53^, and multiple factors such as diet^54^, age^55^, and antibiotic administration^56^ can alter the composition of its microbiome. Our metric predicts that these factors potentially affect the ability of the gut to sustain MGEs and change the contents of mobile elements in gut microbiome. This notion provides new insights on reversing antibiotic resistance in complex microbial communities^57^.

Our results also have implications for the engineering of microbial consortia. Engineering complex bacterial communities in useful ways remains challenging due to the lack of understanding of the ecological principles and intercellular metabolic interactions^58^. Mobile genetic elements are potentially powerful tools for function-oriented microbiota engineering^59^. Our modeling framework and the MGE persistence potential can guide such efforts. In particular, the persistence potential reveals how a few key kinetic parameters predict the approximate abundance of the MGE.

## Methods

### Strains and plasmids

The compositions of the eight engineered communities are shown in Table S1. *E.coli* strain MG1655 without fluorescence markers was denoted as strain X^16^. *E.coli* strain DA26735 with chromosomal BFP and chloramphenicol resistance (Cm^R^) was denoted as strain B^16,60^, and *E.coli* strain DA32838 with chromosomal dTomato and Cm^R^ was denoted as strain R^16,60^. All three strains carry helper F plasmid F_HR_ that expresses tetracycline resistance (Tet^R^)^16^. F_HR_ is not transmissible but encodes the conjugation machinery that mobilize plasmid K. Plasmid K expresses GFP under the control of a strong constitutive PR promoter, and expresses kanamycin resistance (Kan^R^)^16,61^. Plasmid K also carries *oriT*, so it can be transferred through conjugation^16,60^.

The multi-plasmid community was composed of *E.coli* strain MG1655 and DH5α. These two strains were distinguished from each other via blue-white screening on X-gal plates, where MG1655 and DH5α colonies were blue and white, respectively. These communities transferred five conjugative plasmids: F’ (lncF, Tet^R^), PCU1 (lncN, Amp^R^),R388 (lncW, Tm^R^), R6K (lncX, Strp^R^), and RP4 (lncP, Kan^R^). These five plasmids are compatible with each other and carry different antibiotic resistance markers. The plasmids were distinguished from each other via selective plating.

### Long-term dynamics of the engineered microbial communities

Our methods for measuring the long-term plasmid dynamics are based on the protocols established by Lopatkin *et al.16*. Single colonies of three strains (X, B and R) carrying plasmid F_HR_ and K were grown overnight at 37°C for 16 hours with shaking (250 rpm) in LB culture (LB broth from APEX) containing appropriate antibiotics (100 µg/mL Cm, 50 µg/mL Kan, or 20 µg/mL Tet). The overnight cultures were resuspended in M9 medium (M9CA medium broth powder from Amresco, supplemented with 0.1 mg/mL thiamine, 2 mM MgSO_4_, 0.1 mM CaCl_2_, and 0.4% w/v glucose) without antibiotics, and diluted to the initial density of 10^6^ cells/mL. We constructed seven communities using the combinations of these three strains: (a) X; (b) B; (c) R; (d) X+B; (e) X+R; (f) B+R; (g) X+R+B. For the communities (d)-(g), the members of each community were mixed in the equal ratio, and the mixtures were diluted to the initial density of 10^6^ cells/mL. The cells were then distributed in a 96-well plate to a final volume of 200 µL/well, and each community had 12 replicates. The 96-well plate was covered with an AeraSeal^TM^ film sealant (Sigma-Aldrich, SKU A9224) followed by a Breath-Easy sealing membrane (Sigma-Aldrich, SKU Z380059). Plates were shaken at 250 rpm at 37°C for 23 hours. This was denoted as day 0. On day 1, the 12 replicates of each community were divided into four groups, each with three replicates. From each well, 2 µL was removed for flow cytometry. The four groups were subjected to four dilution ratios (10^3^ ×, 10^4^ ×, 10^5^ ×, 10^6^ ×). The new plates were sealed using both membranes and placed back into the incubator. The same protocols were repeated daily from day 1 to day 15.

For the community transferring multiple plasmids, we first transformed the five plasmids into MG1655, respectively, and obtained five plasmid-carrying strains (M^F’^, M^PCU1^, M^R388^, M^R6K^, and M^RP4^). Single colonies of the six strains (M^F’^, M^PCU1^, M^R388^, M^R6K^, M^RP4^, and DH5α) were grown overnight at 37°C for 16 hours with shaking (250 rpm) in LB culture containing appropriate antibiotics (20 µg/mL Tet, 100 µg/mL Amp, 10 µg/mL Tm, 50 µg/mL Strp, and 50 µg/mL Kan, respectively). The overnight cultures were resuspended in M9 medium without antibiotics, and diluted to the initial density of 10^6^ cells/mL. The cells of these six strains were mixed in a ratio of 1:1:1:1:1:5 (M^F’^: M^PCU1^: M^R388^: M^R6K^: M^RP4^: DH5α), and the mixtures were diluted to the initial density of 10^6^ cells/mL. The cells were then distributed in a 96-well plate to a final volume of 200 µL/well, with 12 replicates. The 96-well plate was covered with an AeraSeal^TM^ film sealant followed by a Breath-Easy sealing membrane. Plates were shaken at 250 rpm at 37°C for 23 hours. This was denoted as day 0. At day 1, the 12 replicates of each community were divided into four groups, each with three replicates. From each well, 10 µL was removed for selective plating. The four groups were subjected to four dilution ratios (10^3^ ×, 10^4^ ×, 10^5^ ×, 10^6^ ×), respectively. The new plates were sealed using both membranes and placed back to the incubator. The same protocols were repeated daily from day 1 to day 15. The ratio between MG1655 and DH5α cells was determined by plating on the X-gal plates (100 µg/mL X-gal, 1mM IPTG). The relative abundance of each plasmid was determined by plating on the plates with the corresponding antibiotics (20 µg/mL Tet, 100 µg/mL Amp, 10 µg/mL Tm, 50 µg/mL Strp, and 50 µg/mL Kan).

### Flow cytometry

The community composition and plasmid abundance were quantified using a flow cytometer (MACSQuant^®^ VYB Analyzer). From day 1 to day 15, the overnight cultures were resuspended and diluted to 1: 1000 in 200 µL fresh M9 media before running through the flow. The channels of emission detectors were set as V1 (450/50 nm) for BFP, Y2 (615/20 nm) for dTomato, and B1 (525/50 nm) for GFP. For each sample, 10,000 cells were collected. All data analysis was performed using FlowJo 10.5.3.

### Measuring the fitness costs and conjugation efficiencies of the mobilizable plasmids

Singles colonies of cells that carry the plasmid K (denoted X^K^, B^K^, and R^K^) or do not carry the plasmid K (denoted X^0^, B^0^ and R^0^) were grown overnight at 37°C for 16 hours with shaking in LB with appropriate antibiotics. The cultures were resuspended in M9 medium without antibiotics and then diluted in 1: 10^+^ ratio. The growth curves of these six cell types were measured using a plate reader (TECAN infinite M200 PRO). Six replicates per cell type were used for quantification. The growth rate constants were calculated as the effective growth rates in exponential phases (e.g., the first 5 hours). First, we smoothed the growth curves to filter out the random noises from the plate reader. We then plotted the increments *ΔN*/*Δt* with regards to the cell density *N*. Linear regression was performed, and the slope was obtained as the growth rate constants. The fitness cost *α* was determined by normalizing the growth rates of plasmid-free populations X^0^, B^0^, and R^0^ by the growth rates of plasmid-carrying populations X^K^, B^K^, and R^K^, respectively. The *λ* values were obtained through *λ* = *α* − 1. The fitness costs of the five conjugative plasmids (F’, PCU1, R388, R6K, and RP4) in the host strain MG1655 were measured in the same way as described.

Conjugation efficiencies were estimated using the protocols established by Lopatkin *et al.* with modifications^16^. X^K^, B^K^, and R^K^ served as the donors, and X^0^, B^0^, and R^0^ served as the transconjugants. To distinguish donors with transconjugants, we transformed the recipients with another non-mobilizable plasmid pJM31, which carried ColE1 origin and expressed ampicillin resistance (Amp^R^). Therefore, donors, recipients, and transconjugants can be distinguished by different resistance markers: Kan for donors, Amp for recipients, and Kan+Amp for transconjugants. Overnight cultures of donors and recipients in LB media with appropriate selection agents were resuspended and diluted (1:100) in fresh LB media. Cells were incubated at 37°C with shaking for 2 hours until they reached exponential phase. The cells were then resuspended in M9 media and mixed in 1:1 ratio with a total volume of 200 µL. Mixtures were incubated at room temperature (25°C) for 1 hour without shaking. The donor, recipient, and transconjugant densities were measured by diluting the mixtures (1:10^6^ for donor and recipient, 1:10^4^ for transconjugant), and spreading three replicates onto corresponding selective plates. The conjugation efficiency was obtained as 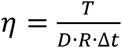, where T, D, R stand for the cell densities of transconjugant, donor and recipient, respectively. Before being plugged into the model, the measured values of *η* need to be normalized with respect to the maximum carrying capacity *N*_*m*_. We estimated *N*_*m*_ to be 6 × 10^8^ cells/mL, which corresponds to OD600≈ 1.2. The dilution rates D were calculated from the daily dilution ratio *ε* by 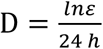^62^. Therefore, dilution ratios of 10^3^ ×, 10^4^ ×, 10^5^ ×, 10^6^ × are equal to dilution rates of 0.2878 *h*^−1^, 0.3838 *h*^−1^, 0.4797 *h*^−1^, and 0.5756 *h*^−1^, respectively. The loss rate *κ* of plasmid K is very small compared with dilution rates D. We used the value (*κ* = 0.001 *h*^−1^) measured by Lopatkin *et al*.^16^.

The conjugation efficiency of the five conjugative plasmids were measured using similar protocols. Since the abundance of DH5α became negligible in the cocultures at the end of the long-term experiments, only the conjugation between MG1655 cells were considered. For plasmid F’ (Tet^R^), PCU1 (Amp^R^),R388 (Tm^R^), and R6K (Strp^R^), MG1655 transformed with the non-mobilizable plasmid K-(Kan^R^, same as the plasmid K but without the transfer origin) served as the recipient. For plasmid RP4 (Kan^R^), MG1655 transformed with the non-mobilizable plasmid pJM31 (Amp^R^) served as the recipient.

## Acknowledgments

We thank Allison J. Lopatkin, Cheemeng Tan, Hong Qian, Terry Hwa, and Lawrence David for thorough reading and comments on an earlier draft of the manuscript and James P. Hall for kindly providing access to the original data of his published work. We thank Ryan Tsoi, Helena Ma, Feilun Wu, Nan Luo, Jia Lu Zach Holmes, Katherine Duncker and Andrea Weiss for comments on the manuscript. We also thank Nicolas Buchler for access to the flow cytometer and Firas Midani for flow cytometry trouble shooting. This work is partially supported by the National Institutes of Health (LY, and R01A1125604 and R01GM110494) and David and Lucile Packard Foundation (LY). The funders had no role in study design, data collection and analysis, decision to publish, or preparation of the manuscript.

## Author contributions

TW conceived the research, designed and performed both modeling and experimental analyses, interpreted the results, and wrote the manuscript. LY conceived the research and assisted in research design, data interpretation, and manuscript writing.

## Data availability

All data is available in the main text and the supplementary information or on request from the corresponding author.

## References

1 Frost, L. S., Leplae, R., Summers, A. O. & Toussaint A. Mobile genetic elements: the agents of open source evolution. Nature Reviews Microbiology 3, 722 (2005).

2 Ochman, H., Lawrence, J. & Groisman E. Lateral gene transfer and the nature of bacterial innovation. Nature 405, 299–304 (2000).

3 Jørgensen, T. S., Xu, Z., Hansen, M. A., Sørensen, S. J. & Hansen L. H. Hundreds of circular novel plasmids and DNA elements identified in a rat cecum metamobilome. PloS one 9, e87924 (2014).

4 Kav, A. B. et al. Insights into the bovine rumen plasmidome. Proceedings of the National Academy of Sciences 109, 5452–5457 (2012).

5 Zhang, T., Zhang, X.-X. & Ye L. Plasmid metagenome reveals high levels of antibiotic resistance genes and mobile genetic elements in activated sludge. PloS one 6, e26041 (2011).

6 Dahlberg, C., Linberg, C., Torsvik, V. L. & Hermansson M. Conjugative plasmids isolated from bacteria in marine environments show various degrees of homology to each other and are not closely related to well-characterized plasmids. Appl. Environ. Microbiol. 63, 4692–4697 (1997).

7 Brito, I. L. et al. Mobile genes in the human microbiome are structured from global to individual scales. Nature 535, 435 (2016).

8 Dennis, J. J. The evolution of IncP catabolic plasmids. Current Opinion in Biotechnology 16, 291–298 (2005).

9 Pilla, G. & Tang C. M. Going around in circles: virulence plasmids in enteric pathogens. Nature Reviews Microbiology 16, 484–495 (2018).

10 Mercer, R. et al. Genetic determinants of heat resistance in Escherichia coli. Frontiers in microbiology 6, 932 (2015).

11 Broaders, E., O’Brien, C., Gahan, C. G. & Marchesi J. R. Evidence for plasmid-mediated salt tolerance in the human gut microbiome and potential mechanisms. FEMS microbiology ecology 92 (2016).

12 Stokes, H. W. & Gillings M. R. Gene flow, mobile genetic elements and the recruitment of antibiotic resistance genes into Gram-negative pathogens. FEMS microbiology reviews 35, 790–819 (2011).

13 Ferreiro, A., Crook, N., Gasparrini, A. J. & Dantas G. Multiscale evolutionary dynamics of host-associated microbiomes. Cell 172, 1216–1227 (2018).

14 Iwasaki, W. & Takagi T. Rapid pathway evolution facilitated by horizontal gene transfers across prokaryotic lineages. PLoS genetics 5, e1000402 (2009).

15 Heuer, H. & Smalla K. Plasmids foster diversification and adaptation of bacterial populations in soil. FEMS microbiology reviews 36, 1083–1104 (2012).

16 Lopatkin, A. J. et al. Persistence and reversal of plasmid-mediated antibiotic resistance. Nature communications 8, 1689 (2017).

17 Johnsen, P. J. et al. Factors affecting the reversal of antimicrobial-drug resistance. The Lancet Infectious Diseases 9, 357–364 (2009).

18 Berg, P., Baltimore, D., Brenner, S., Roblin, R. O. & Singer M. F. Asilomar conference on recombinant DNA molecules. Science 188, 991–994 (1975).

19 Wright, O., Delmans, M., Stan, G.-B. & Ellis T. GeneGuard: a modular plasmid system designed for biosafety. ACS synthetic biology 4, 307–316 (2015).

20 Thiry, M. & Cingolani D. Optimizing scale-up fermentation processes. TRENDS in Biotechnology 20, 103–105 (2002).

21 Silva, F., Queiroz, J. A. & Domingues F. C. Plasmid DNA fermentation strategies: influence on plasmid stability and cell physiology. Applied microbiology and biotechnology 93, 2571–2580 (2012).

22 Slater, F. R., Bailey, M. J., Tett, A. J. & Turner S. L. Progress towards understanding the fate of plasmids in bacterial communities. FEMS Microbiology Ecology 66, 3–13 (2008).

23 Sørensen, S. J., Bailey, M., Hansen, L. H., Kroer, N. & Wuertz S. Studying plasmid horizontal transfer in situ: a critical review. Nature Reviews Microbiology 3, 700 (2005).

24 Bergstrom, C. T., Lipsitch, M. & Levin B. R. Natural selection, infectious transfer and the existence conditions for bacterial plasmids. Genetics 155, 1505–1519 (2000).

25 Stewart, F. M. & Levin B. R. The population biology of bacterial plasmids: a priori conditions for the existence of conjugationally transmitted factors. Genetics 87, 209–228 (1977).

26 Condit, R., Stewart, F. M. & Levin B. R. The population biology of bacterial transposons: a priori conditions for maintenance as parasitic DNA. The American Naturalist 132, 129–147 (1988).

27 Chao, L., Levin, B. R. & Stewart, F. M. A complex community in a simple habitat: an experimental study with bacteria and phage. Ecology 58, 369–378 (1977).

28 Leclerc, Q. J., Lindsay, J. A. & Knight G. M. Mathematical modelling to study the horizontal transfer of antimicrobial resistance genes in bacteria: current state of the field and recommendations. Journal of the Royal Society Interface 16, 20190260 (2019).

29 Lozupone, C. A., Stombaugh, J. I., Gordon, J. I., Jansson, J. K. & Knight R. Diversity, stability and resilience of the human gut microbiota. Nature 489, 220 (2012).

30 Curtis, T. P., Sloan, W. T. & Scannell J. W. Estimating prokaryotic diversity and its limits. Proceedings of the National Academy of Sciences 99, 10494–10499 (2002).

31 Munck, C., Sheth, R. U., Freedberg, D. E. & Wang H. H. Recording mobile DNA in the gut microbiota using an Escherichia coli CRISPR-Cas spacer acquisition platform. Nature Communications 11, 1–11 (2020).

32 Martin-Serrano, J. & Neil S. J. Host factors involved in retroviral budding and release. Nature Reviews Microbiology 9, 519–531 (2011).

33 Lenski, R. E. & Levin B. R. Constraints on the coevolution of bacteria and virulent phage: a model, some experiments, and predictions for natural communities. The American Naturalist 125, 585–602 (1985).

34 Levin, B. R., Stewart, F. M. & Chao L. Resource-limited growth, competition, and predation: a model and experimental studies with bacteria and bacteriophage. The American Naturalist 111, 3–24 (1977).

35 Condit, R. The evolution of transposable elements: conditions for establishment in bacterial populations. Evolution 44, 347–359 (1990).

36 Hall, J. P., Wood, A. J., Harrison, E. & Brockhurst M. A. Source–sink plasmid transfer dynamics maintain gene mobility in soil bacterial communities. Proceedings of the National Academy of Sciences 113, 8260–8265 (2016).

37 Loftie-Eaton, W. et al. Compensatory mutations improve general permissiveness to antibiotic resistance plasmids. Nature ecology & evolution 1, 1354 (2017).

38 Hall, J. P., Williams, D., Paterson, S., Harrison, E. & Brockhurst M. A. Positive selection inhibits gene mobilization and transfer in soil bacterial communities. Nature ecology & evolution 1, 1348 (2017).

39 Harrison, E., Guymer, D., Spiers, A. J., Paterson, S. & Brockhurst M. A. Parallel compensatory evolution stabilizes plasmids across the parasitism-mutualism continuum. Current Biology 25, 2034–2039 (2015).

40 Kottara, A., Hall, J. P., Harrison, E. & Brockhurst M. A. Variable plasmid fitness effects and mobile genetic element dynamics across Pseudomonas species. FEMS microbiology ecology 94, fix172 (2017).

41 Dahlberg, C. & Chao L. Amelioration of the cost of conjugative plasmid carriage in Eschericha coli K12. Genetics 165, 1641–1649 (2003).

42 Fischer, E. A. et al. The IncI1 plasmid carrying the bla CTX-M-1 gene persists in in vitro culture of a Escherichia coli strain from broilers. BMC microbiology 14, 77 (2014).

43 Porse, A., Schønning, K., Munck, C. & Sommer M. O. Survival and evolution of a large multidrug resistance plasmid in new clinical bacterial hosts. Molecular biology and evolution 33, 2860–2873 (2016).

44 Bohannan, B. J. & Lenski R. E. Effect of resource enrichment on a chemostat community of bacteria and bacteriophage. Ecology 78, 2303–2315 (1997).

45 Jones, G. W., Baines, L. & Genthner F. J. Heterotrophic bacteria of the freshwater neuston and their ability to act as plasmid recipients under nutrient deprived conditions. Microbial ecology 22, 15–25 (1991).

46 Winter, S. et al. Host-Derived Nitrate Boosts Growth of E. coli in the Inflamed Gut. Science 339, 708–711 (2013).

47 Stecher, B. et al. Gut inflammation can boost horizontal gene transfer between pathogenic and commensal Enterobacteriaceae. Proceedings of the National Academy of Sciences 109, 1269–1274 (2012).

48 Wotzka, S. Y., Nguyen, B. D. & Hardt, W.-D. Salmonella Typhimurium diarrhea reveals basic principles of enteropathogen infection and disease-promoted DNA exchange. Cell host & microbe 21, 443–454 (2017).

49 Lopatkin, A. J. et al. Antibiotics as a selective driver for conjugation dynamics. Nature microbiology 1, 16044 (2016).

50 Korem, T. et al. Growth dynamics of gut microbiota in health and disease inferred from single metagenomic samples. Science 349, 1101–1106 (2015).

51 Arnoldini, M., Cremer, J. & Hwa T. Bacterial growth, flow, and mixing shape human gut microbiota density and composition. Gut microbes 9, 559–566 (2018).

52 Dionisio, F., Matic, I., Radman, M., Rodrigues, O. R. & Taddei F. Plasmids spread very fast in heterogeneous bacterial communities. Genetics 162, 1525–1532 (2002).

53 Mitreva, M. & Consortium, H. M. P. Structure, function and diversity of the healthy human microbiome. Nature 486, 207–214 (2012).

54 David, L. A. et al. Diet rapidly and reproducibly alters the human gut microbiome. Nature 505, 559 (2014).

55 Yatsunenko, T. et al. Human gut microbiome viewed across age and geography. Nature 486, 222–227 (2012).

56 Willing, B. P., Russell, S. L. & Finlay B. B. Shifting the balance: antibiotic effects on host–microbiota mutualism. Nature Reviews Microbiology 9, 233–243 (2011).

57 Andersson, D. I. & Hughes D. Antibiotic resistance and its cost: is it possible to reverse resistance? Nature Reviews Microbiology 8, 260 (2010).

58 Sheth, R. U., Cabral, V., Chen, S. P. & Wang H. H. Manipulating bacterial communities by in situ microbiome engineering. Trends in Genetics 32, 189–200 (2016).

59 Ronda, C., Chen, S. P., Cabral, V., Yaung, S. J. & Wang H. H. Metagenomic engineering of the mammalian gut microbiome in situ. Nature methods 16, 167–170 (2019).

60 Gullberg, E. et al. Selection of resistant bacteria at very low antibiotic concentrations. PLoS pathogens 7 (2011).

61 Dimitriu, T. et al. Genetic information transfer promotes cooperation in bacteria. Proceedings of the National Academy of Sciences 111, 11103–11108 (2014).

62 Balagaddé, F. K., You, L., Hansen, C. L., Arnold, F. H. & Quake S. R. Long-term monitoring of bacteria undergoing programmed population control in a microchemostat. Science 309, 137–140 (2005).

