## supplementary information for "The Persistence Potential of Mobile Genetic Elements"

**This PDF file includes:**

Supplementary Methods
Supplementary Text
Figs. S1 to S5
Tables S1 to S20

**Section 1: Supplementary methods**

**Strains and plasmids**

The compositions of the eight engineered communities are shown in Table S1. *E.coli* strain MG1655 without fluorescence markers was denoted as strain X. *E.coli* strain DA26735 with chromosomal BFP and chloramphenicol resistance ($Cm^R$) was denoted as strain B, and *E.coli* strain DA32838 with chromosomal dTomato and $Cm^R$ was denoted as strain R. All three strains carry helper F plasmid $F_{HR}$ that expresses tetracycline resistance ($Tet^R$). $F_{HR}$ is not transmissible but encodes the conjugation machinery that mobilize plasmid K. Plasmid K expresses GFP under the control of a strong constitutive PR promoter, and expresses kanamycin resistance ($Kan^R$). Plasmid K also carries *oriT*, so it can be transferred through conjugation.

The multi-plasmid community was composed of *E.coli* strain MG1655 and DH5α. These two strains were distinguished from each other via blue-white screening on X-gal plates, where MG1655 and DH5α colonies were blue and white, respectively. These communities transferred five conjugative plasmids: F' (lncF, $Tet^R$), PCU1 (lncN, $Amp^R$),R388 (lncW, $Tm^R$), R6K (lncX, $Strp^R$), and RP4 (lncP, $Kan^R$). These five plasmids are compatible with each other and carry different antibiotic resistance markers. The plasmids were distinguished from each other via selective plating.

**Table S1** The strain compositions and plasmids in the eight engineered communities

| communities | strains | plasmids |
|---|---|---|
| 1 | X (MG1655+$F_{HR}$) | K ($Kan^R$, GFP) |
| 2 | B (DA26735+$F_{HR}$) | |
| 3 | R (DA32838+$F_{HR}$) | |
| 4 | X, B | |
| 5 | X, R | |
| 6 | B, R | |
| 7 | X, B, R | |
| 8 | MG1655, DH5$\alpha$ | F' (lncF, $Tet^R$), PCU1 (lncN, $Amp^R$),R388 (lncW, $Tm^R$), R6K (lncX, $Strp^R$), and RP4 (lncP, $Kan^R$) |

### Supplementary Text

**Section 2: Model and Theoretical Analysis**

**2.1 Model development of MGE-centric framework (MCF)**

**2.1.1 An illustrative example: community of two species and two plasmids**

To explain the key concepts of the MGE-centric framework, we will first focus on the plasmid dynamics. For a community of two species and two plasmids, let $s_1$ and $s_2$ represent the abundances of species 1 and 2, respectively. Let $p_{11}$ represent the abundance of species-1 cells that carry plasmid 1, and $p_{12}$ represent the abundance of species-1 cells that carry plasmid 2. In a similar way, we can define $p_{21}$ and $p_{22}$. MCF describes how the community composition ($s_i$, $i = 1,2$) and plasmid distribution ($p_{ij}$, $i = 1,2$ and $j = 1,2$) change with time. First, we assume all plasmids are compatible with each other, which means that they can coexist in the same host cell. The dynamics of this community can then be described by six ODEs

$$\frac{ds_1}{dt} = \alpha_1 \mu_1^e s_1 - D s_1, \quad [1]$$

$$\frac{ds_2}{dt} = \alpha_2 \mu_2^e s_2 - D s_2, \quad [2]$$

$$\frac{dp_{11}}{dt} = \beta_{11}\mu_{11}^e p_{11} + (s_1 - p_{11})(\eta_{111}p_{11} + \eta_{121}p_{21}) - (\kappa_{11} + D)p_{11}, \quad [3]$$

$$\frac{dp_{12}}{dt} = \beta_{12}\mu_{12}^e p_{12} + (s_1 - p_{12})(\eta_{211}p_{12} + \eta_{221}p_{22}) - (\kappa_{12} + D)p_{12}, \quad [4]$$

$$\frac{dp_{21}}{dt} = \beta_{21}\mu_{21}^e p_{21} + (s_2 - p_{21})(\eta_{122}p_{21} + \eta_{112}p_{11}) - (\kappa_{21} + D)p_{21}, \quad [5]$$

$$\frac{dp_{22}}{dt} = \beta_{22}\mu_{22}^e p_{22} + (s_2 - p_{22})(\eta_{222}p_{22} + \eta_{212}p_{12}) - (\kappa_{22} + D)p_{22}, \quad [6]$$

These ODEs constitute the main part of MCF. $s_i$ increases by cell division, the effective growth rate of which is represented by $\mu_i^e$. Each plasmid might cause a fitness burden or benefit on the growth of the host cell, and $\alpha_i$ represents the combined growth effect of all the plasmids that species $i$ carries. $D$ is the dilution rate. $p_{ij}$ increases by cell division ($\mu_{ij}^e$) or horizontal transfer through conjugation ($\eta_{jki}$). The effective growth rate of the host cells that carry plasmid $j$ is represented by $\mu_{ij}^e$, which will be smaller than $\mu_i^e$ if the plasmid is burdensome and larger than $\mu_i^e$ if the plasmid brings benefit. The combined effect of all other plasmids on the division rates of $p_{ij}$ is described by $\beta_{ij}$. Plasmid $j$ can also be transferred horizontally from species $k$ to species $i$ at a rate constant of $\eta_{jki}$. Therefore, the influx of $p_{ij}$ from species $k$ is obtained as $(s_i - p_{ij})\eta_{jki}p_{kj}$. The rate constant of plasmid loss is represented by $\kappa_{ij}$.

$\mu_i^e$ represents the effective growth rate of the 'empty' cells (cells that do not carry any plasmids), while $\mu_{ij}^e$ is the effective growth rate of the cells that are only equipped with the plasmid $j$. The effective growth rates $\mu_i^e$ and $\mu_{ij}^e$ are calculated from the maximum growth rates (represented by $\mu_i$ and $\mu_{ij}$, respectively) and the carrying capacity $c_i$ through $\mu_i^e = \mu_i c_i$, $\mu_{ij}^e = \mu_{ij} c_i$. $\mu_{ij}$ is linked to $\mu_i$ through the fitness cost of $p_{ij}$, which is denoted $\lambda_{ij}$. We assume that the maximum growth rate is inversely proportional to the fitness cost. Therefore, the relationship between $\mu_{ij}$ and $\mu_i$ is obtained as $\mu_{ij} = \mu_i/(1 + \lambda_{ij})$. The plasmid is burdensome with positive $\lambda_{ij}$ and beneficial with negative $\lambda_{ij}$. To obtain the combined cost of all the plasmids that $s_i$ carries, we calculated the weighted average of their costs as $\overline{\lambda_i} = \frac{p_{i1}}{s_i}\lambda_{i1} + \frac{p_{i2}}{s_i}\lambda_{i2}$. Then, the formulation of $\alpha_i$ can be obtained as $\alpha_i = \frac{1}{1+\overline{\lambda_i}}$, which leads to

$$\alpha_1 = \frac{s_1}{s_1 + p_{11}\lambda_{11} + p_{12}\lambda_{12}}, \quad [7]$$

$$\alpha_2 = \frac{s_2}{s_2 + p_{21}\lambda_{21} + p_{22}\lambda_{22}}. \quad [8]$$

Formulating $\beta_{ij}$, however, requires prior information about the distribution patterns of the other plasmids in $p_{ij}$ cells. For a general understanding, we equilibrated its distribution in $p_{ij}$ cells as its distribution in $s_i$ cells. Then, similar to the definition of $\alpha_i$, $\beta_{ij}$ can be obtained as

$$\beta_{11} = \frac{s_1(1+\lambda_{11})}{s_1(1+\lambda_{11}) + p_{12}\lambda_{12}}, \quad [9]$$

$$\beta_{12} = \frac{s_1(1+\lambda_{12})}{s_1(1+\lambda_{12}) + p_{11}\lambda_{11}}, \quad [10]$$

$$\beta_{21} = \frac{s_2(1+\lambda_{21})}{s_2(1+\lambda_{21}) + p_{22}\lambda_{22}}, \quad [11]$$

$$\beta_{22} = \frac{s_2(1+\lambda_{22})}{s_2(1+\lambda_{22}) + p_{21}\lambda_{21}}. \quad [12]$$

Here, we assume the two species compete with each other and follow the logistic growth. Therefore, the growth capacities can be formulated as $c_1 = c_2 = 1 - s_1 - s_2$.

##### 2.1.2 The matrix form of the MGE-centric framework

Using vectors and matrices, our MGE-centric framework can be summarized into two equations

$$\frac{dS}{dt} = A \circ U_S \circ S \circ C - D \cdot S; \quad [13]$$

$$\frac{dP}{dt} = B \circ U_P \circ P \circ \bar{C} + (\bar{S} - P) \circ H \circ \bar{P} - D \cdot P - K \circ P. \quad [14]$$

Here, we still use communities of two species and two plasmids to illustrate the formations of each term in the equations. The formulations of communities with higher numbers of species and plasmids can be obtained in exactly the same way. In the equations, '∘' is the Hadamard product. $S$ and $P$ represent the vectors or matrices of the abundances of species and plasmids, respectively:

$$S = \begin{bmatrix} s_1 \\ s_2 \end{bmatrix}, P = \begin{bmatrix} p_{11} & p_{12} \\ p_{21} & p_{22} \end{bmatrix}.$$

$A$, $U_S$ and $C$ are the vector forms of $\alpha_i$, $\mu_i$ and $c_i$, respectively:

$$A = \begin{bmatrix} \alpha_1 \\ \alpha_2 \end{bmatrix}, U_S = \begin{bmatrix} \mu_1 \\ \mu_2 \end{bmatrix}, C = \begin{bmatrix} c_1 \\ c_2 \end{bmatrix}.$$

$B$, $U_P$ and $\bar{C}$ are the matrix forms of $\beta_{ij}$, $\mu_{ij}$ and $c_i$, respectively:

$$B = \begin{bmatrix} \beta_{11} & \beta_{12} \\ \beta_{21} & \beta_{22} \end{bmatrix}, U_P = \begin{bmatrix} \mu_{11} & \mu_{12} \\ \mu_{21} & \mu_{22} \end{bmatrix}, \bar{C} = \begin{bmatrix} c_1 & c_1 \\ c_2 & c_2 \end{bmatrix}.$$

$\bar{S}$ is the expanded matrix of the vector $S$,

$$\bar{S} = \begin{bmatrix} s_1 & s_1 \\ s_2 & s_2 \end{bmatrix}.$$

$H$contains the horizontal transfer rates, with the form

$$H = \begin{bmatrix} [\eta_{111} & \eta_{121}] & [\eta_{211} & \eta_{221}] \\ [\eta_{112} & \eta_{122}] & [\eta_{212} & \eta_{222}] \end{bmatrix},$$

where $\eta_{jki}$ is the conjugation rate constant of plasmid $j$ from species $k$ to species $j$. $\bar{P}$ is expressed as

$$\bar{P} = \begin{bmatrix} \begin{bmatrix} p_{11} \\ p_{21} \end{bmatrix} & \begin{bmatrix} p_{12} \\ p_{22} \end{bmatrix} \\ \begin{bmatrix} p_{11} \\ p_{21} \end{bmatrix} & \begin{bmatrix} p_{12} \\ p_{22} \end{bmatrix} \end{bmatrix}.$$

$K$ contains the plasmid loss rates

$$K = \begin{bmatrix} \kappa_{11} & \kappa_{12} \\ \kappa_{21} & \kappa_{22} \end{bmatrix}.$$

$\kappa_{ij}$ represents the loss rate of plasmid $j$ in species $i$.

##### 2.1.3 Generalization of MGE-centric framework

The basic principles of the MCF have been discussed in section 2.1.1 and 2.1.2. Here, we will generalize the framework to communities with more species and plasmids. First, we discuss communities composed of $m$ species and $n$ plasmids. Let $s_i$ ($i = 1,2,\dots,m$) represent the abundance of species $i$, and $p_{ij}$ ($i = 1,2,\dots,m$ and $j = 1,2,\dots,n$) represent the abundance of plasmid $j$-carrying cells in species $i$. We assume all MGEs are compatible with each other. The dynamics of this community can then be described as follows:

$$\frac{ds_i}{dt} = \alpha_i \mu_i s_i c_i - D s_i, \quad [15]$$

$$\frac{dp_{ij}}{dt} = \beta_{ij}\mu_{ij}p_{ij}c_i + \left(s_i - p_{ij}\right)\sum_{k=1}^{m} \eta_{jki}p_{kj} - \left(\kappa_{ij} + D\right)p_{ij}. \quad [16]$$

The definition of each parameter is shown in section 2.1.1. The formulation of $\alpha_i$ in this general case becomes

$$\alpha_i = \frac{s_i}{s_i + \sum_{j=1}^{n}(p_{ij}\lambda_{ij})}, \quad [17]$$

and $\beta_{ij}$ becomes

$$\beta_{ij} = \frac{s_i\left(1+\lambda_{ij}\right)}{s_i\left(1+\lambda_{ij}\right) + \sum_{\{k:\, k\neq j\}}(p_{ik}\lambda_{ik})}. \quad [18]$$

Next, we consider plasmid incompatibility. Let $G_l$ represent the incompatibility groups ($l = 1,2,\dots,t$, and $t \leq n$) and $Z_j$ represent the map from plasmid index $j$ to the index of its incompatibility group $l$. For instance, for plasmid $j$ and $k$, if $Z_j = Z_k$, these two plasmids belong to the same incompatibility group and will not coexist stably in the same host cell. Therefore, the modified ODEs for $p_{ij}$ become

$$\frac{dp_{ij}}{dt} = \beta_{ij}\mu_{ij}p_{ij}c_i + \left(s_i - \sum_{\{h:\, Z_h = Z_j\}} p_{ih}\right)\sum_{k=1}^{m} \eta_{jki}p_{kj} - \left(\kappa_{ij} + D\right)p_{ij}, \quad [19]$$

$$\beta_{ij} = \frac{s_i(1+\lambda_{ij})}{s_i(1+\lambda_{ij}) + \sum_{\{h:\, Z_h \neq Z_j\}}(p_{ih}\lambda_{ih})}. \quad [20]$$

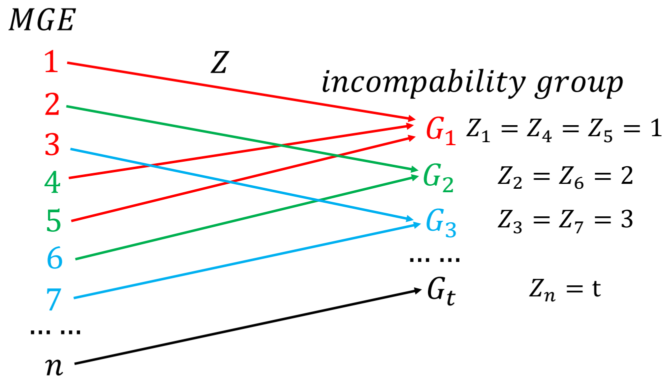


An example of a community containing 200 species and two incompatible plasmids is shown in Fig. S1C.

Finally, we discuss the general formulation of the growth capacities $c_i$. We take niche partitioning and species interactions into consideration. The general form of $c_i$ is expressed as follows

$$c_i = e_i - \sum_{k=1}^{m} x_{ki} s_k, \quad [21]$$

where $e_i$ is the maximum carrying capacity of the niche that species $i$ locates, and $x_{ki}$ represents the strength of the interaction from species $k$ to species $i$. In its simplest case, we applied the logistic growth model, which means that we only considered the competition among species. Similar to the way in which we characterize plasmid incompatibility, we distributed the $m$ species into $r$ niches $W_1$, $W_2$,…, $W_r$ ($r \leq m$). The maximum carrying capacity of these niches are $y_1$, $y_2$,…, $y_r$, respectively. Let $V_i$ represent the map from the species index $i$ to its niche index. Therefore, $e_i = y_{V_i}$. Then, the formulation of $C_i$ that we used in our analysis was obtained as

$$c_i = y_{V_i} - \sum_{\{k:\, V_k = V_i\}} s_k. \quad [22]$$

The ODEs, and the formulations of $\alpha$, $\beta$ and $c$, constitute the body of our framework. Now we are ready for numerical simulations.

**2.2 Analysis on the computation complexities of subpopulation-centric framework (SCF) and MGE-centric framework (MCF)**

We consider a microbial community of $m$ species and $n$ mobile elements. In SCF, the population can be divided into $m \cdot 2^n$ subpopulations. Each subpopulation represents a unique combination of species and MGEs. Thus, SCF requires $m \cdot 2^n$ ODEs. However, MCF contains only $m(n+1)$ ODEs, $m$ of which describe population dynamics, and $mn$ of which describe MGE dynamics.

The computation complexities of these two frameworks can also be quantified by the number of parameters. In SCF, each subpopulation has its unique growth rate. Thus, $m \cdot 2^n$ growth rates are required. For each of the $n$ MGEs, there are $m \cdot 2^{n-1}$ subpopulations that carry it and another $m \cdot 2^{n-1}$ subpopulations that do not carry it. The total number of transfer rates can then be obtained as $n(m \cdot 2^{n-1})^2 = nm^2 \cdot 2^{2n-2}$. Every subpopulation that carries the specific mobile element has a unique rate of losing it. Therefore, the number of MGE loss rates in SCF is $mn \cdot 2^{n-1}$.

In MCF, because we consider the growth rates of only the MGE-free cells as well as the fitness cost of each mobile element on the MGE-free populations, only $m(n+1)$ growth rate constants are needed. For horizontal transfer, we only consider the transfer rates of a specific MGE across different species, so the total number of transfer rates is also greatly reduced to $nm^2$. In a similar way, for each mobile element, we only consider its loss in each species instead of in each subpopulation, so the number of loss rates are reduced to $mn$. The results of this computation complexity analysis are summarized in Table S2.

**Table S2** Analysis of the computation complexities of SCF and MCF

| | | SCF | MCF |
|---|---|---|---|
| Number of ODEs | | $m \cdot 2^n$ | $m(n+1)$ |
| Number of parameters | Growth rates | $m \cdot 2^n$ | $m(n+1)$ |
| | Segregation loss rates | $mn \cdot 2^{n-1}$ | $mn$ |
| | Dilution rates | 1 | 1 |
| | Conjugation efficiencies | $nm^2 \cdot 2^{2n-2}$. | $nm^2$ |

#### 2.3 Discrepancy between SCF and MCF

For microbial communities with multiple species and one mobile element, we compared the performances of the two frameworks using numerical simulations. We assigned the same sets of randomized parameters to both models. The relative abundances of MGEs, quantified as the fractions of MGE-carrying cells at steady states, from the two models were aligned with each other (Fig. S2A). From 1000 repeated numerical tests, no qualitative discrepancy was detected. The quantitative discrepancy shows dependence on fitness cost. Lower average growth burden or benefit gives rise to better match.

For communities with multiple mobile elements coexisting, SCF requires more parameters to predict the MGE dynamics (Table S1). Here we focus on a community with one species and two MGEs (denoted A and B, respectively). Let $\lambda_A$ and $\lambda_B$ represent the individual fitness costs when the mobile elements exist separately. In MCF, the combined fitness cost of A and B when they coexist in the same host cell can be derived from $\lambda_A$ and $\lambda_B$. However, in SCF, one additional parameter $\lambda_{AB}$ is required to characterize the combined effect. We calculated the outputs under different values of $\lambda_A$ and $\lambda_B$ as well as $\lambda_{AB}$ (Fig. S2B). The results suggest that the two models match well over a wide range of fitness costs.

#### 2.4 The application of the framework to lytic bacteriophage and transposons

##### 2.4.1 Lytic bacteriophages

During infection, lytic bacteriophages replicate in the host cells. After a short latent time, the infected host cells lyse, releasing the resident phages. Because the host cannot grow in the lytic cycle, our framework requires modifications to accommodate lytic phage dynamics. The new ODEs are shown below:

$$\frac{ds}{dt} = \mu s(1 - s) - \eta sp - Ds, \quad [23]$$

$$\frac{dp}{dt} = \beta\eta sp - Dp - \kappa p. \quad [24]$$

$s$ and $p$ stand for the densities of cells and phages, respectively. $\mu$ is the growth rate constant of cells. $\eta$ is the rate constant at which phages encounter and irreversibly infect the cells. One infected cell releases $\beta$ phages after lysing. $D$ and $q$ represent the dilution and phage loss rates, respectively.

This model represents a simplified version of the classic one[33]. The condition for phage persistence ($q > 0$) can be derived as

$$\beta\eta > \frac{\mu}{\mu - D}(D + \kappa). \quad [25]$$

Therefore, we defined the persistence potential in this case as

$$\omega = \frac{\beta\eta}{\frac{\mu}{\mu - D}(D + \kappa)}. \quad [26]$$

For heterogeneous communities, the formulation of $\omega$ can be generalized as described in Section 2.4.

##### 2.4.2 Transposons

Transposons are the genetic elements that locate on chromosomes or plasmids. The dynamical processes of transposons are more complex than those of plasmids. The transfer of transposons is mediated by plasmid conjugation, and the intercellular transfer of the transposon includes two steps: (1) the transposition of the transposon from chromosome to plasmid and (2) the conjugation of the plasmid across different cells[26].

For a population in which all cells carry the conjugative plasmid, let $s_i$ $(i = 1,2, \dots, m)$ represent the abundance of the species $i$, and $p_{ij}$ $(i = 1,2, \dots, m$ and $j = 1,2, \dots, n)$ represent the abundance of transposon $j$-carrying cells in the species $i$. The overall transfer rate of the transposon is represented by $\eta$. $\lambda_{ij}$ stands for the fitness cost of the $j$th transposon on the $i$th species. The transposon can be lost through plasmid segregation loss and excision from chromosome or plasmid[35]. The rate constant of transposon loss is represented by $\kappa$. The transposon dynamics can be described by the same ODEs described in Section 2.1.3.

#### 2.5 Development of MGE persistence potential

We started with a microbial community of $m$ species and one mobile element. Assuming that the community is homogeneous, i.e. all the species carry the same kinetic parameters, the dynamics of this community can be described by $2m$ ODEs:

$$\frac{ds_i}{dt} = \frac{s_i}{s_i + \lambda p_i}\mu s_i\left(\frac{1}{m} - s_i\right) - Ds_i, \quad [27]$$

$$\frac{dp_i}{dt} = \frac{\mu}{1+\lambda} p_i \left(\frac{1}{m} - s_i\right) + m\eta(s_i - p_i)p_i - (D+\kappa)p_i, \quad [28]$$

$$(i = 1,2,\ldots,m).$$

We then assumed that all $s_i$ or $p_i$ start with the same initial densities ($s_1^0 = s_2^0 = \cdots = s_m^0$ and $p_1^0 = p_2^0 = \cdots = p_m^0$). Therefore, at any time points, $s_1 = s_2 = \cdots = s_m$ and $p_1 = p_2 = \cdots = p_m$. Thus, this system can be simplified as two ODEs:

$$\frac{dx}{dt} = \frac{x}{x+\lambda y} \mu_e x(1-x) - Dx, \quad [29]$$

$$\frac{dy}{dt} = \frac{\mu_e}{1+\lambda} y(1-x) + \eta(x-y)y - (D+\kappa)y, \quad [30]$$

where $x = ms_i$, $y = mp_i$ and $\mu_e = \frac{\mu}{m}$. At steady state, with $x \neq 0$ and $y \neq 0$, Eq. 29 can be transformed to

$$y = \frac{-\mu_e x^2 + (\mu_e - D)x}{D\lambda}, \quad [31]$$

and Eq. 30 can be transformed to

$$y = \left(1 - \frac{\mu_e}{\alpha\eta}\right)x + \left(\frac{\mu_e}{\alpha\eta} - \frac{D+\kappa}{\eta}\right). \quad [32]$$

To derive the criterion of MGE persistence in this simplified case, we then sought the conditions under which Eq. 31 and Eq. 32 give rise to solutions with positive and feasible values of $x$ and y ($0 < \mathrm{x} \leq 1, 0 < \mathrm{y} \leq \mathrm{x}$). Mathematically, this is a quadratic problem, and the criterion can be analytically derived out as follows

$$\eta > \frac{\mu}{\mu - mD} \cdot \frac{\alpha(D+\kappa) - D}{\alpha}. \quad [33]$$

We defined the MGE persistence potential as

$$\omega = \frac{\eta}{\frac{\mu}{\mu - mD} \cdot \frac{\alpha(D+\kappa) - D}{\alpha}}. \quad [34]$$

Therefore, this criterion can be expressed as $\omega > 1$.

Next, we sought to generalize the criterion to heterogeneous communities with multiple mobile elements coexisting. We reformulated $\omega_j$ (the $\omega$ value of MGE $j$) in the following manner:

$$\omega_j = \frac{\overline{\eta_j}}{\frac{\bar{\mu}}{\bar{\mu} - \sigma D}\left(D + \overline{\kappa_j} - \frac{D}{1+\overline{\lambda_j}}\right)}, \quad [35]$$

where $\bar{\mu}$, $\overline{\kappa_j}$, $\overline{\lambda_j}$ and $\overline{\eta_j}$ are the weighted average values of the corresponding parameters with regards to the population sizes. $\sigma$ characterizes the diversity of the community and is calculated by $\sigma = e^{-\sum_{i=1}^{m} \frac{s_i}{s_T} ln \frac{s_i}{s_T}}$, where $s_T$ represents the total size of the community ($s_T = s_1 + s_1 + \cdots +$

$s_m$ ). $\bar{\mu}$ , $\bar{\kappa_J}$ ,and $\bar{\lambda_J}$ are defined as follows: $\bar{\mu} = \frac{1}{s_T}\sum_{k=1}^{m}\mu_k s_k$ , $\bar{\kappa_J} = \frac{1}{s_T}\sum_{k=1}^{m}\kappa_{jk}s_k$ , $\bar{\lambda_J} = \frac{1}{s_T}\sum_{k=1}^{m}\lambda_{jk}s_k$. When formulating the overall conjugation efficiency $\bar{\eta_J}$, however, the contributions of both donor and recipient population sizes should be considered. Thus $\bar{\eta_J}$ is defined as: $\bar{\eta_J} = \frac{1}{s_T^2}\sum_{ik=1,2,\ldots,m}\eta_{jik}s_i s_k$.

#### Section 3: Analysis of Literature Data

Here, we summarized the literature data from 13 previous studies. The microbial communities analyzed in these studies covered one, two, or three populations, and up to three MGEs. The data source of these data are shown in Table S3.

**Table S3** The referred papers and the data sources

| References | Data sources |
|---|---|
| Hall JP, Wood AJ, Harrison E, Brockhurst MA. Source–sink plasmid transfer dynamics maintain gene mobility in soil bacterial communities. Proceedings of the National Academy of Sciences. 2016 Jul 19;113(29):8260-5. | 1. The plasmid abundance data, which were visualized in Fig. 1A and C of this referred paper, was obtained from DRYAD (https://datadryad.org/bitstream/handle/10255/dryad.119190/Figure1Data.csv?sequence=1)<br>2. The parameter values were obtained from Table S1 in the Supporting Information of this referred paper. |
| Lopatkin AJ, Meredith HR, Srimani JK, Pfeiffer C, Durrett R, You L. Persistence and reversal of plasmid-mediated antibiotic resistance. Nature communications. 2017 Nov 22;8(1):1689. | 1. The parameter values were obtained from Supplementary Tables 2 and 3 and Supplementary Figures 3 and 6 of this referred paper.<br>2. The plasmid abundance data were extracted from Figures 2, 3, and 4 of this referred paper. |
| Loftie-Eaton W, Bashford K, Quinn H, Dong K, Millstein J, Hunter S, Thomason MK, Merrikh H, Ponciano JM, Top EM. Compensatory mutations improve general permissiveness to antibiotic resistance plasmids. Nature ecology & evolution. 2017 Sep;1(9):1354. | 1. The plasmid abundance data, which were visualized in Fig. 1 of this referred paper, was obtained from DRYAD (https://datadryad.org/bitstream/handle/10255/dryad.147859/ResultsSection2.1.csv?sequence=1)<br>2. The parameter values were extracted from Fig. 3 and Fig. S2 of this referred paper. |
| Hall JP, Williams D, Paterson S, Harrison E, Brockhurst MA. Positive selection inhibits gene mobilization and transfer in soil bacterial communities. Nature ecology & evolution. 2017 Sep;1(9):1348. | 1.The parameter values were obtained from Table S1 in the Supporting Information of 'Hall JP, Wood AJ, Harrison E, Brockhurst MA. Source–sink plasmid transfer dynamics maintain gene mobility in soil bacterial communities. Proceedings of the National Academy of Sciences. 2016 Jul 19;113(29):8260-5.'<br>2.The plasmid abundance data, which were visualized in Fig. 3b of this referred paper, was obtained from DRYAD |

| | |
|---|---|
| | (https://datadryad.org/bitstream/handle/10255/dryad.148630/S4_Tn5042_PF_Data.csv?sequence=1) |
| Harrison E, Guymer D, Spiers AJ, Paterson S, Brockhurst MA. Parallel compensatory evolution stabilizes plasmids across the parasitism-mutualism continuum. Current Biology. 2015 Aug 3;25(15):2034-9. | 1. The parameter values were extracted from Figure 1 and Figure S1 of this referred article.<br>2. The plasmid abundance data were extracted from Figure S2 of the Supplemental Information of this referred article. |
| Kottara A, Hall JP, Harrison E, Brockhurst MA. Variable plasmid fitness effects and mobile genetic element dynamics across Pseudomonas species. FEMS microbiology ecology. 2017 Dec 4;94(1):fix172. | 1. The selection rates were extracted from Figure 1 of the referred article.<br>2. The plasmid abundance data were extracted from Figure 2 of the referred article. |
| Dahlberg C, Chao L. Amelioration of the cost of conjugative plasmid carriage in Escherichia coli K12. Genetics. 2003 Dec 1;165(4):1641-9. | The fitness cost of evolved combinations, the prevalence of evolved three R1 populations, and the plasmid transfer rates of evolved R1 plasmids were obtained from Table 2, 3 and 4 of the referred article, respectively. |
| Fischer EA, Dierikx CM, van Essen-Zandbergen A, van Roermund HJ, Mevius DJ, Stegeman A, Klinkenberg D. The IncI1 plasmid carrying the *bla*$_{CTX\text{-}M\text{-}1}$ gene persists in in vitro culture of an *Escherichia coli* strain from broilers. BMC microbiology. 2014 Dec;14(1):77. | 1. The parameter values were extracted from Tables 1, 2, and 3 as well as the main text of this inferred article.<br>2. The plasmid abundance data were extracted from Fig. 2 and 3 of this referred article. |
| Porse A, Schønning K, Munck C, Sommer MO. Survival and evolution of a large multidrug resistance plasmid in new clinical bacterial hosts. Molecular biology and evolution. 2016 Aug 8;33(11):2860-73. | 1. The parameters were obtained from supplementary Tables S3 and S4 of this referred paper.<br>2. The plasmid abundance data were abstracted from Fig. 4 of this referred paper. |
| Levin BR, Stewart FM, Chao L. Resource-limited growth, competition, and predation: a model and experimental studies with bacteria and bacteriophage. The American Naturalist. 1977 Jan 1;111(977):3-24. | 1. The parameters were obtained from the main text of this referred paper.<br>2. The abundance data of bacteriophage T2 were abstracted from Fig. 5 and Fig. 8 of this referred paper. |
| Lenski RE, Levin BR. Constraints on the coevolution of | 1. The parameters were obtained from the Table 1 of this referred paper. |

| | |
|---|---|
| bacteria and virulent phage: a model, some experiments, and predictions for natural communities. The American Naturalist. 1985 Apr 1;125(4):585-602. | 2. The abundance data of bacteriophage T4 were abstracted from Fig. 1 of this referred paper. |
| Bohannan BJ, Lenski RE. Effect of resource enrichment on a chemostat community of bacteria and bacteriophage. Ecology. 1997 Dec 1;78(8):2303-15. | 1. The parameters were obtained from Lenski *et al*. 1985 (*2*).<br>2. The abundance data of bacteriophage T4 were abstracted from Fig. 3 of this referred paper. |
| Condit R. The evolution of transposable elements: conditions for establishment in bacterial populations. Evolution. 1990 Mar;44(2):347-59. | 1. The parameters were obtained from Table 1 and 2 of this referred paper.<br>2. The abundance data of the transposons were abstracted from Fig. 1 and 2 of this referred paper. |

**3.1 Source-sink plasmid transfer dynamics maintain gene mobility in soil bacterial communities**

Hall JP *et al*. investigated how the presence of alternative host species affects plasmid population dynamics. They cultured *Pseudomonas fluorescens* SBM25 and *Pseudomonas putica* KT2440 either individually or together. The populations started with 50% frequency of a mercury resistance plasmid, pQBR57. The samples were transferred into fresh media either with or without mercuric chloride selections. The dynamics of the bacteria populations were tracked over 65 transfers.

To verify the predictive power of MGE persistence potential, we calculated the $\omega$ values and the corresponding relative abundance of plasmid pQBR57 in each of their experiments. Because the parameter values under mercuric chloride treatment were lacking, we only focused on their experimental data without mercuric selections. Additionally, we only considered the data after transfer 29 once the systems were close to steady states. With these two constraints, the original data we reused in this paper are summarized in Table S4. The parameter values are summarized in Table S5. The $\lambda$ factors in our model can be calculated from the $\alpha$ values shown in Table S5 by $\lambda_1 = \frac{1}{\alpha_1} - 1$, $\lambda_2 = \frac{1}{\alpha_2} - 1$. The conjugation efficiency was measured using the end-point method, with units of $cell^{-1}hr^{-1}$, and was dependent on the cell density. In our model, we used the cell density-independent conjugation rate constant $\eta$, by normalizing the measured values with respect to the maximum carrying capacities $N_m$. For the experiments with single species, $N_m$ values are shown in Table S5. For coculture experiments, we estimated $N_m$ as the mean value of total population size. In this work, cell death instead of dilution was considered. Since both death rate $d$ and dilution rate $D$ occupy the same position in our equations, the persistence potential can be calculated as $\omega = \frac{\bar{\eta}}{\frac{\bar{\mu}}{\bar{\mu}-\sigma d}\left(d+\bar{\kappa}-\frac{d}{1+\bar{\lambda}}\right)}$. The pQBR57 relative abundance was calculated as the ratio of plasmid-carrying cells to the total population size. The $\omega$ values and the corresponding abundances are summarized in Table S6.

**Table S4**. The original data of the relative abundance of plasmid pQBR57 from 'Source-sink plasmid transfer dynamics maintain gene mobility in soil bacterial communities.'

| | | single species | | | | coculture | | | |
|---|---|---|---|---|---|---|---|---|---|
| | | *Pseudomonas fluorescens* | | *Pseudomonas putica* | | *Pseudomonas fluorescens* | | *Pseudomonas putica* | |
| | | Cell density | Plasmid density | Cell density | Plasmid density | Cell density | Plasmid density | Cell density | Plasmid density |
| Transfer 29 | A | $7.1 \times 10^8$ | $6.9 \times 10^8$ | $2.4 \times 10^8$ | 0 | $1 \times 10^9$ | 0 | $1.2 \times 10^8$ | 0 |
| | B | $5 \times 10^8$ | $5 \times 10^8$ | $1.6 \times 10^8$ | 0 | $5 \times 10^8$ | 0 | $5 \times 10^7$ | 0 |
| | C | $6.1 \times 10^8$ | $4.2 \times 10^8$ | $1.2 \times 10^8$ | 0 | $6 \times 10^8$ | $3.7 \times 10^8$ | $1.5 \times 10^8$ | 0 |
| | D | $9.1 \times 10^8$ | $2.8 \times 10^7$ | $5.4 \times 10^7$ | 0 | $8.8 \times 10^8$ | 0 | $9.5 \times 10^7$ | 0 |
| | E | $5.2 \times 10^8$ | $1.6 \times 10^7$ | $1.1 \times 10^8$ | 0 | $5.5 \times 10^8$ | $5.4 \times 10^8$ | $5.3 \times 10^7$ | $1.7 \times 10^6$ |
| | F | $4.9 \times 10^8$ | $3.2 \times 10^8$ | $1.2 \times 10^8$ | 0 | $6.4 \times 10^8$ | $2 \times 10^7$ | $5 \times 10^7$ | $4.7 \times 10^6$ |
| Transfer 35 | A | $6 \times 10^8$ | $6 \times 10^8$ | $9 \times 10^7$ | 0 | $3.6 \times 10^8$ | 0 | $1.4 \times 10^7$ | 0 |
| | B | $4.4 \times 10^8$ | $4 \times 10^8$ | $8.7 \times 10^7$ | 0 | $4.8 \times 10^8$ | $4.5 \times 10^7$ | $4.2 \times 10^7$ | 0 |
| | C | $6.2 \times 10^8$ | $4.5 \times 10^8$ | $1.3 \times 10^8$ | 0 | $3.9 \times 10^8$ | $3.9 \times 10^8$ | $3.1 \times 10^7$ | 0 |
| | D | $3.6 \times 10^8$ | $3 \times 10^8$ | $9.8 \times 10^7$ | $3.1 \times 10^6$ | $4.5 \times 10^8$ | 0 | $7 \times 10^7$ | 0 |
| | E | $5 \times 10^8$ | $5 \times 10^8$ | $6.8 \times 10^7$ | 0 | $3.2 \times 10^8$ | $3.2 \times 10^8$ | $7 \times 10^7$ | $1.8 \times 10^7$ |
| | F | $5.1 \times 10^8$ | $4.1 \times 10^8$ | $8.6 \times 10^7$ | 0 | $3.4 \times 10^8$ | $1.7 \times 10^8$ | $2.5 \times 10^7$ | $7.9 \times 10^5$ |
| Transfer 41 | A | $4.9 \times 10^8$ | $4.9 \times 10^8$ | $5.6 \times 10^7$ | 0 | $4.8 \times 10^8$ | 0 | $5.6 \times 10^7$ | 0 |
| | B | $4.8 \times 10^8$ | $4.7 \times 10^8$ | $8.8 \times 10^7$ | 0 | $2.3 \times 10^8$ | $2 \times 10^8$ | $2.5 \times 10^7$ | $3.2 \times 10^6$ |
| | C | $3.7 \times 10^8$ | $3.4 \times 10^8$ | $1.3 \times 10^8$ | 0 | $3.4 \times 10^8$ | $3.3 \times 10^8$ | $6.4 \times 10^7$ | $2 \times 10^6$ |
| | D | $4.6 \times 10^8$ | $4.6 \times 10^8$ | $1.4 \times 10^8$ | 0 | $4.1 \times 10^8$ | $3 \times 10^8$ | $4.5 \times 10^7$ | $2.8 \times 10^6$ |
| | E | $4.6 \times 10^8$ | $4.6 \times 10^8$ | $9.2 \times 10^7$ | 0 | $5 \times 10^8$ | $4.8 \times 10^8$ | $7 \times 10^7$ | $3.5 \times 10^7$ |
| | F | $3.9 \times 10^8$ | $3.8 \times 10^8$ | $7.4 \times 10^7$ | 0 | $4.8 \times 10^8$ | $4.5 \times 10^8$ | $3.9 \times 10^7$ | $3.7 \times 10^6$ |

| | | | | | | | | | |
|---|---|---|---|---|---|---|---|---|---|
| Transfer 47 | A | $5.74 \times 10^8$ | $5.74 \times 10^8$ | $1.4 \times 10^8$ | 0 | $4.9 \times 10^8$ | 0 | $5.3 \times 10^7$ | 0 |
| | B | $6.8 \times 10^8$ | $6.6 \times 10^8$ | $1.4 \times 10^8$ | 0 | $6.7 \times 10^8$ | $4 \times 10^8$ | $7.3 \times 10^7$ | $2.3 \times 10^6$ |
| | C | $4.9 \times 10^8$ | $4.9 \times 10^8$ | $1.2 \times 10^8$ | 0 | $4.4 \times 10^8$ | $4.4 \times 10^8$ | $3.9 \times 10^7$ | 0 |
| | D | $5.04 \times 10^8$ | $5.04 \times 10^8$ | $1.4 \times 10^7$ | 0 | $5.3 \times 10^8$ | $3.6 \times 10^8$ | $4.2 \times 10^7$ | $1.3 \times 10^6$ |
| | E | $6 \times 10^8$ | $6 \times 10^8$ | $1.1 \times 10^8$ | 0 | $5.9 \times 10^8$ | $5.9 \times 10^8$ | $9.5 \times 10^7$ | $3 \times 10^7$ |
| | F | $6.2 \times 10^8$ | $6 \times 10^8$ | $1.2 \times 10^7$ | 0 | $6.2 \times 10^8$ | $4.7 \times 10^8$ | $2.8 \times 10^7$ | $1.8 \times 10^6$ |
| Transfer 53 | A | $6.2 \times 10^8$ | $5.2 \times 10^8$ | $1.1 \times 10^8$ | $3.4 \times 10^6$ | $5.3 \times 10^8$ | 0 | $6.9 \times 10^7$ | 0 |
| | B | $4.8 \times 10^8$ | $4.3 \times 10^8$ | $1.1 \times 10^8$ | 0 | $4.1 \times 10^8$ | $3.8 \times 10^8$ | $6.6 \times 10^7$ | 0 |
| | C | $4.5 \times 10^8$ | $4.2 \times 10^8$ | $1.2 \times 10^8$ | 0 | $4.6 \times 10^8$ | $2.9 \times 10^8$ | $1.5 \times 10^8$ | $9.1 \times 10^6$ |
| | D | $5 \times 10^8$ | $5 \times 10^8$ | $8.6 \times 10^7$ | 0 | $3.4 \times 10^8$ | $2.1 \times 10^8$ | $4 \times 10^7$ | $2.5 \times 10^6$ |
| | E | $4.8 \times 10^8$ | $4.8 \times 10^8$ | $1 \times 10^8$ | 0 | $3.6 \times 10^8$ | $3.6 \times 10^8$ | $6.9 \times 10^7$ | $2.2 \times 10^7$ |
| | F | $5 \times 10^8$ | $4.7 \times 10^8$ | $9.4 \times 10^7$ | 0 | $4.4 \times 10^8$ | $4 \times 10^8$ | $3.6 \times 10^7$ | $2.3 \times 10^6$ |
| Transfer 59 | A | $4.7 \times 10^8$ | $4.7 \times 10^8$ | $5.2 \times 10^7$ | 0 | $5.8 \times 10^8$ | 0 | $5.9 \times 10^7$ | 0 |
| | B | $4.7 \times 10^8$ | $3.9 \times 10^8$ | $9.1 \times 10^7$ | 0 | $4.9 \times 10^8$ | $4.8 \times 10^8$ | $3.6 \times 10^7$ | $2.3 \times 10^6$ |
| | C | $5.1 \times 10^8$ | $5 \times 10^8$ | $1.1 \times 10^8$ | 0 | $4.2 \times 10^8$ | $2.6 \times 10^8$ | $6.3 \times 10^7$ | 0 |
| | D | $3.7 \times 10^8$ | $3.7 \times 10^8$ | $3.9 \times 10^7$ | 0 | $4.3 \times 10^8$ | $3.5 \times 10^8$ | $5.6 \times 10^7$ | $1.8 \times 10^6$ |
| | E | $4 \times 10^8$ | $4 \times 10^8$ | $8.1 \times 10^7$ | 0 | $3.3 \times 10^8$ | $3.3 \times 10^8$ | $3.6 \times 10^7$ | 0 |
| | F | $3.9 \times 10^8$ | $3.7 \times 10^8$ | $1 \times 10^8$ | 0 | $4 \times 10^8$ | $2.7 \times 10^8$ | $8.3 \times 10^7$ | 0 |
| Transfer 65 | A | $3.2 \times 10^8$ | $3.2 \times 10^8$ | $1.3 \times 10^8$ | 0 | $2.9 \times 10^8$ | 0 | $1.7 \times 10^8$ | 0 |
| | B | $7.9 \times 10^8$ | $7.2 \times 10^8$ | $1.6 \times 10^8$ | 0 | $2.6 \times 10^8$ | $2 \times 10^8$ | $1.1 \times 10^8$ | 0 |
| | C | $5.4 \times 10^8$ | $4.9 \times 10^8$ | $1.3 \times 10^8$ | 0 | $5.4 \times 10^8$ | $1.9 \times 10^8$ | $1.1 \times 10^8$ | $3.4 \times 10^6$ |
| | D | $3.6 \times 10^8$ | $3.1 \times 10^8$ | $1.5 \times 10^8$ | 0 | $3.4 \times 10^8$ | $1.4 \times 10^8$ | $9.6 \times 10^7$ | 0 |

| | | | | | | | | | |
|---|---|---|---|---|---|---|---|---|---|
| | E | $6.5 \times 10^8$ | $6.3 \times 10^8$ | $1.2 \times 10^8$ | 0 | $4.4 \times 10^8$ | $4.4 \times 10^8$ | $1.8 \times 10^8$ | $2.3 \times 10^7$ |
| | F | $6.3 \times 10^8$ | $3.1 \times 10^8$ | $1.2 \times 10^8$ | 0 | $3.6 \times 10^8$ | $9 \times 10^7$ | $8.6 \times 10^7$ | 0 |

* The plasmid abundance data, which were visualized in Fig. 1A and C of this referred paper, were obtained from DRYAD (https://datadryad.org/bitstream/handle/10255/dryad.119190/Figure1Data.csv?sequence=1).

**Table S5** The parameter values used in 'Source-sink plasmid transfer dynamics maintain gene mobility in soil bacterial communities'

| Name | Variable measured | Estimate |
|---|---|---|
| $\mu_1$ | *Pseudomonas fluorescens* growth rate | $0.091\ h^{-1}$ |
| $\mu_2$ | *Pseudomonas putica* growth rate | $0.186\ h^{-1}$ |
| $\alpha_1$ | Effect of plasmid carriage on *Pseudomonas fluorescens* growth rate | 0.73 |
| $\alpha_2$ | Effect of plasmid carriage on *Pseudomonas putica* growth rate | 0.85 |
| $\eta_{11}$ | Intraspecific conjugation rate in *Pseudomonas fluorescens* | $10^{-11}\ cell^{-1} \cdot h^{-1}$ |
| $\eta_{12}$ | Interspecific conjugation rate from *Pseudomonas fluorescens* to *Pseudomonas putica* | $10^{-14}\ cell^{-1} \cdot h^{-1}$ |
| $\eta_{21}$ | Interspecific conjugation rate from *Pseudomonas putica* to *Pseudomonas fluorescens* | $10^{-11.5}\ cell^{-1} \cdot h^{-1}$ |
| $\eta_{22}$ | Intraspecific conjugation rate in *Pseudomonas putica* | $10^{-14}\ cell^{-1} \cdot h^{-1}$ |
| $\kappa$ | *Segregation rate* | $1 \times 10^{-4}\ h^{-1}$ |
| $N_{m1}$ | Carrying capacity of *Pseudomonas fluorescens* | $6.01 \times 10^8$ |
| $N_{m2}$ | Carrying capacity of *Pseudomonas putica* | $1.1 \times 10^8$ |
| $d$ | Death rate | $0.009\ h^{-1}$ |

* The parameter values were obtained from Table S1 in the Supporting Information of this referred paper. We changed the letters of each parameter to fit the denotations of our model.

**Table S6** Plasmid pQBR57 persistence potential and the corresponding relative abundance in 'Source-sink plasmid transfer dynamics maintain gene mobility in soil bacterial communities'

| | | | Plasmid persistence potential | Plasmid relative abundance |
|---|---|---|---|---|
| Transfer 29 | Single species | *Pseudomonas fluorescens* | 2.14 | $0.56 \pm 0.43$ |
| | | *Pseudomonas putica* | $7.22 \times 10^{-4}$ | 0 |
| | Coculture | | $1.39 \pm 0.08$ | $0.24 \pm 0.37$ |
| Transfer 35 | Single species | *Pseudomonas fluorescens* | 2.14 | $0.88 \pm 0.11$ |
| | | *Pseudomonas putica* | $7.22 \times 10^{-4}$ | $0.005 \pm 0.013$ |
| | Coculture | | $1.41 \pm 0.09$ | $0.39 \pm 0.43$ |

| | | | | |
|---|---|---|---|---|
| Transfer 41 | Single species | *Pseudomonas fluorescens* | 2.14 | $0.98 \pm 0.03$ |
| | | *Pseudomonas putica* | $7.22 \times 10^{-4}$ | 0 |
| | Coculture | | $1.39 \pm 0.05$ | $0.68 \pm 0.34$ |
| Transfer 47 | Single species | *Pseudomonas fluorescens* | 2.14 | $0.99 \pm 0.02$ |
| | | *Pseudomonas putica* | $7.22 \times 10^{-4}$ | 0 |
| | Coculture | | $1.42 \pm 0.06$ | $0.62 \pm 0.34$ |
| Transfer 53 | Single species | *Pseudomonas fluorescens* | 2.14 | $0.96 \pm 0.04$ |
| | | *Pseudomonas putica* | $7.22 \times 10^{-4}$ | $0.005 \pm 0.013$ |
| | Coculture | | $1.36 \pm 0.10$ | $0.59 \pm 0.33$ |
| Transfer 59 | Single species | *Pseudomonas fluorescens* | 2.14 | $0.96 \pm 0.06$ |
| | | *Pseudomonas putica* | $7.22 \times 10^{-4}$ | 0 |
| | Coculture | | $1.38 \pm 0.06$ | $0.61 \pm 0.34$ |
| Transfer 65 | Single species | *Pseudomonas fluorescens* | 2.14 | $0.86 \pm 0.19$ |
| | | *Pseudomonas putica* | $7.22 \times 10^{-4}$ | 0 |
| | Coculture | | $1.13 \pm 0.13$ | $0.35 \pm 0.26$ |

##### 3.2 Persistence and reversal of plasmid-mediated antibiotic resistance

Lopatkin AJ *et al.* studied the conjugation-assisted persistence of different plasmids. Here we focused on three groups of experiments presented in their paper.

(1) They introduced a mobilizable plasmid (denoted plasmid K) into the *E.coli* strain MG1655 (denoted strain B) equipped with a helper plasmid $F_{HR}$. $F_{HR}$ is not self-transmissible but encodes the conjugation machinery and helps the transfer of plasmid K (kanamycin-resistant). They then quantified the long-term dynamics of plasmid K abundance. Different antibiotic concentrations were applied to change the growth effects of plasmid K. Another mobilizable plasmid denoted C (Chloramphenicol-resistant) that carries mCherry marker was also tested in strain B. To demonstrate conjugation-assisted persistence of natural plasmids, they also quantified the dynamics of plasmids 168, 193, R388, 41, RP4, PCU1, and R6K in an *E.coli* MG1655 strain with chromosomally integrated dTomato (denoted as strain R).

(2) To reverse the conjugation-assisted persistence, they used linoleic acid to inhibit conjugation and phenothiazine to enhance the plasmid segregation error. This treatment significantly reduced plasmid persistence.

(3) They tested plasmid persistence in complex communities: (a) strain B mixed with strain R, carrying plasmid K; (b) strain B, carrying plasmids K and C; (c) strain B, strain R, and strain Y (*E.coli* MG1655 with YFP), carrying plasmids R6K, RP4, and R388. In experiments (a) and (b), different concentrations of kanamycin or chloramphenicol were added to change the fitness cost of the plasmids.

Lopatkin *et al.* also investigated the persistence of a non-mobilizable plasmid K-, which is identical to plasmid K except that it does not carry *oriT* and therefore cannot be transferred through conjugation.

The kinetic parameters of these experiments are summarized in Table S7. The $\alpha$ values represent the magnitude of plasmid burdens. The $\lambda$ factors in our model can be calculated from the $\alpha$ values through $\lambda = \alpha - 1$. To transform the measured values of conjugation efficiency $\eta$ to the values used in their mathematical model, Lopatkin *et al.* normalized $\eta$ respect to the maximum carrying capacity $N_m = 1 \times 10^9\ cell \cdot mL^{-1}$. However, in their experiments, $\eta$ was measured using cells in stationary phase, so they accounted for the physiological influence on $\eta$ by multiplying the measured conjugation efficiency by a factor of $2.5 \times 10^3$. In this way, we calculated the plasmid persistence potential in these experiments, as shown in Table S8. For the non-mobilizable plasmid $K^-$, the plasmid persistence potential $\omega$ is 0 because $\eta$ equals 0. When the value of $\frac{\bar{\mu}}{\bar{\mu}-\sigma D}\left(D + \bar{\kappa} - \frac{D}{1+\bar{\lambda}}\right)$ is positive, the plasmid persistence potential is described as $+0$, and when $\frac{\bar{\mu}}{\bar{\mu}-\sigma D}\left(D + \bar{\kappa} - \frac{D}{1+\bar{\lambda}}\right)$ is negative, the plasmid persistence potential is described as $-0$.

**Table S7** Parameter values used in 'Persistence and reversal of plasmid-mediated antibiotic resistance'

| Name | Variable measured | Estimate |
|---|---|---|
| $\mu_B^K$ | Growth rate of $B^K$ (or $B^{K-}$) population | $0.3\ h^{-1}$ |
| $\mu_R^K$ | Growth rate of $R^K$ population | $0.29\ h^{-1}$ |
| $\mu_B^C$ | Growth rate of $B^C$ population | $0.28\ h^{-1}$ |
| $\mu_R^{41}$ | Growth rate of $R^{41}$ population | $0.14\ h^{-1}$ |
| $\mu_R^{168}$ | Growth rate of $R^{168}$ population | $0.19\ h^{-1}$ |
| $\mu_R^{193}$ | Growth rate of $R^{193}$ population | $0.21\ h^{-1}$ |
| $\mu_R^{RP4}$ | Growth rate of $R^{RP4}$ population | $0.20\ h^{-1}$ |
| $\mu_R^{R6K}$ | Growth rate of $R^{R6K}$ population | $0.25\ h^{-1}$ |
| $\mu_R^{PCU1}$ | Growth rate of $R^{PCU1}$ population | $0.11\ h^{-1}$ |
| $\mu_R^{R388}$ | Growth rate of $R^{PCU1}$ population | $0.28\ h^{-1}$ |
| $\alpha_B^K$ | Growth burden of plasmid K (or K–) on strain B | 1.02 |
| $\alpha_B^{K'}$ | Growth burden of plasmid K (or K–) on strain B with 0.5 $\mu g/ml$ Kanamycin treatment | 0.97 |
| $\alpha_B^{K''}$ | Growth burden of plasmid K (or K–) on strain B with 2 $\mu g/ml$ Kanamycin treatment | 0.42 |
| $\alpha_B^{K'''}$ | Growth burden of plasmid K on strain B with 0.5 $\mu g/ml$ Chloramphenicol treatment | 1.035 |
| $\alpha_B^{K''''}$ | Growth burden of plasmid K on strain B with 2 $\mu g/ml$ Chloramphenicol treatment | 1.00 |
| $\alpha_R^K$ | Growth burden of plasmid K (or K–) on strain R | 1.07 |
| $\alpha_R^{K'}$ | Growth burden of plasmid K (or K–) on strain R with 0.5 $\mu g/ml$ Kanamycin treatment | 0.99 |
| $\alpha_R^{K''}$ | Growth burden of plasmid K (or K–) on strain R with 2 $\mu g/ml$ Kanamycin treatment | 0.86 |
| $\alpha_B^C$ | Growth burden of plasmid C on strain B | 1.21 |
| $\alpha_B^{C'}$ | Growth burden of plasmid C on strain B with 0.5 $\mu g/ml$ Chloramphenicol treatment | 0.89 |

| | | |
|---|---|---|
| $\alpha_B^{C''}$ | Growth burden of plasmid C on strain B with $2\,\mu g/ml$ Chloramphenicol treatment | 0.33 |
| $\alpha_B^{C'''}$ | Growth burden of plasmid C on strain B with $0.5\,\mu g/ml$ Kanamycin treatment | 1.5 |
| $\alpha_B^{C''''}$ | Growth burden of plasmid C on strain B with $2\,\mu g/ml$ Kanamycin treatment | 1.00 |
| $\alpha_R^{C}$ | Growth burden of plasmid C on strain R | 1.22 |
| $\alpha_R^{41}$ | Growth burden of plasmid 41 on strain R | 1.37 |
| $\alpha_R^{168}$ | Growth burden of plasmid 168 on strain R | 0.97 |
| $\alpha_R^{193}$ | Growth burden of plasmid 193 on strain R | 0.90 |
| $\alpha_R^{RP4}$ | Growth burden of plasmid RP4 on strain R | 0.90 |
| $\alpha_B^{RP4}$ | Growth burden of plasmid RP4 on strain B | 0.89 |
| $\alpha_Y^{RP4}$ | Growth burden of plasmid RP4 on strain Y | 0.90 |
| $\alpha_R^{R6K}$ | Growth burden of plasmid R6K on strain R | 0.75 |
| $\alpha_B^{R6K}$ | Growth burden of plasmid R6K on strain B | 0.87 |
| $\alpha_Y^{R6K}$ | Growth burden of plasmid R6K on strain Y | 0.92 |
| $\alpha_R^{PCU1}$ | Growth burden of plasmid PCU1 on strain R | 2.80 |
| $\alpha_R^{R388}$ | Growth burden of plasmid R388 on strain R | 0.66 |
| $\alpha_B^{R388}$ | Growth burden of plasmid R388 on strain B | 0.84 |
| $\alpha_Y^{R388}$ | Growth burden of plasmid R388 on strain Y | 0.86 |
| $\eta_K$ | Conjugation efficiency of plasmid K without inhibition | $2.5 \times 10^{-15}\ cell^{-1}h^{-1}mL$ |
| $\eta_C$ | Conjugation efficiency of plasmid C without inhibition | $1.09 \times 10^{-14}\ cell^{-1}h^{-1}mL$ |
| $\eta_{41}$ | Conjugation efficiency of plasmid 41 without inhibition | $1.80 \times 10^{-14}\ cell^{-1}h^{-1}mL$ |
| $\eta_{168}$ | Conjugation efficiency of plasmid 168 without inhibition | $5.94 \times 10^{-15}\ cell^{-1}h^{-1}mL$ |
| $\eta_{193}$ | Conjugation efficiency of plasmid 193 without inhibition | $2.38 \times 10^{-15}\ cell^{-1}h^{-1}mL$ |
| $\eta_{RP4}$ | Conjugation efficiency of plasmid RP4 without inhibition | $1.76 \times 10^{-12}\ cell^{-1}h^{-1}mL$ |
| $\eta_{R6K}$ | Conjugation efficiency of plasmid R6K without inhibition | $9.19 \times 10^{-14}\ cell^{-1}h^{-1}mL$ |
| $\eta_{PCU1}$ | Conjugation efficiency of plasmid PCU1 without inhibition | $1.45 \times 10^{-13}\ cell^{-1}h^{-1}mL$ |
| $\eta_{R388}$ | Conjugation efficiency of plasmid R388 without inhibition | $1.94 \times 10^{-12}\ cell^{-1}h^{-1}mL$ |
| $D$ | Dilution rate | $0.05\ h^{-1}$ |
| $\kappa$ | Segregation loss rate without phenothiazine | $5.2 \times 10^{-4}\ h^{-1}$ |
| $\sigma_K$ | Fold-decrease of conjugation efficiency of plasmid K with inhibition | 2.96 |
| $\sigma_C$ | Fold-decrease of conjugation efficiency of plasmid C with inhibition | 1.54 |

| | | |
|---|---|---|
| $\sigma_{41}$ | Fold-decrease of conjugation efficiency of plasmid 41 with inhibition | 12.43 |
| $\sigma_{168}$ | Fold-decrease of conjugation efficiency of plasmid 168 with inhibition | 53.42 |
| $\sigma_{193}$ | Fold-decrease of conjugation efficiency of plasmid 193 with inhibition | 3.31 |
| $\sigma_{RP4}$ | Fold-decrease of conjugation efficiency of plasmid RP4 with inhibition | 13.06 |
| $\sigma_{R6K}$ | Fold-decrease of conjugation efficiency of plasmid R6K with inhibition | 2.46 |
| $\sigma_{PCU1}$ | Fold-decrease of conjugation efficiency of plasmid PCU1 with inhibition | 13.09 |
| $\sigma_{R388}$ | Fold-decrease of conjugation efficiency of plasmid PCU1 with inhibition | 2.49 |
| $\kappa_{Ph}$ | Segregation loss rate with phenothiazine | $2.1 \times 10^{-3}\ h^{-1}$ |

*The parameter values were obtained from Supplementary Table 2, 3, Supplementary Figure 3 and 6 of this referred paper. We changed the letters of each parameters to fit the denotations of in our model.

**Table S8** Persistence potential and the corresponding relative abundance of plasmid RP4 in 'Persistence and reversal of plasmid-mediated antibiotic resistance'

| | Plasmid persistence potential $\omega$ | Plasmid relative abundance | Description |
|---|---|---|---|
| Simple communities without linoleic acid and phenothiazine | 1.38 | $1 \pm 0.00$ | Plasmid K in strain R |
| | 2.53 | $0.96 \pm 0.07$ | Plasmid C in strain B (selective plating) |
| | 2.37 | $1.02 \pm 0.06$ | Plasmid 41 in strain R |
| | $-10.54$ | $0.91 \pm 0.07$ | Plasmid 168 in strain R |
| | $-0.87$ | $1.01 \pm 0.06$ | Plasmid 193 in strain R |
| | $-625$ | $0.97 \pm 0.21$ | Plasmid RP4 in strain R |
| | $-10.44$ | $0.91 \pm 0.04$ | Plasmid R6K in strain R |
| | $-140.85$ | $0.90 \pm 0.04$ | Plasmid R388 in strain R |
| Simple communities | 0.33 | $0 \pm 0.00$ | Plasmid K in strain R |
| | 1.40 | $0.28 \pm 0.03$ | Plasmid C in strain B |
| | 0.17 | $0 \pm 0.00$ | Plasmid 41 in strain R |

| | | | |
|---|---|---|---|
| with linoleic acid and phenothiazine | 0.38 | 0.02 ± 0.00 | Plasmid 168 in strain R |
| | −0.38 | 0.43 ± 0.03 | Plasmid 193 in strain R |
| | −69.93 | 0.66 ± 0.12 | Plasmid RP4 in strain R |
| | −4.69 | 0.60 ± 0.08 | Plasmid R6K in strain R |
| | 0.68 | 0 ± 0.00 | Plasmid PCU1 in strain R |
| | −59.88 | 0.81 ± 0.04 | Plasmid R388 in strain R |
| Simple communities with antibiotic treatment | 3.48 | 1.00 ± 0.00 | Plasmid K in strain B with 0 $\mu g/ml$ Kanamycin |
| | −5.04 | 1.00 ± 0.01 | Plasmid K in strain B with 0.5 $\mu g/ml$ Kanamycin |
| | −0.06 | 1.00 ± 0.00 | Plasmid K in strain B with 2 $\mu g/ml$ Kanamycin |
| | 2.53 | 0.69 ± 0.16 | Plasmid C in strain B with 0 $\mu g/ml$ Chloramphenicol (flow cytometry) |
| | −3.85 | 0.90 ± 0.04 | Plasmid C in strain B with 0.5 $\mu g/ml$ Chloramphenicol |
| | −0.12 | 0.98 ± 0.02 | Plasmid C in strain B with 2 $\mu g/ml$ Chloramphenicol |
| Complex communities | 1.96 | 0.98 ± 0.04 | Plasmid K in communities $B^K$+$R^K$ with 0 $\mu g/ml$ Kanamycin |
| | −10.33 | 0.99 ± 0.05 | Plasmid K in communities $B^K$+$R^K$ with 0.5 $\mu g/ml$ Kanamycin |
| | −0.17 | 0.97 ± 0.02 | Plasmid K in communities $B^K$+$R^K$ with 2 $\mu g/ml$ Kanamycin |
| | 3.48 | 0.95 ± 0.06 | Plasmid K in communities $B^K$+$B^C$+$B^{CK}$ with 0 $\mu g/ml$ Kanamycin |
| | −5.04 | 0.97 ± 0.06 | Plasmid K in communities $B^K$+$B^C$+$B^{CK}$ with0.5 $\mu g/ml$ Kanamycin |
| | −0.06 | 0.97 ± 0.02 | Plasmid K in communities $B^K$+$B^C$+$B^{CK}$ with 2 $\mu g/ml$ Kanamycin |
| | 2.37 | 0.91 ± 0.15 | Plasmid K in communities $B^K$+$B^C$+$B^{CK}$ with 0.5 $\mu g/ml$ Chloramphenicol |
| | 10.02 | 0.82 ± 0.07 | Plasmid K in communities $B^K$+$B^C$+$B^{CK}$ with2 $\mu g/ml$ Chloramphenicol |
| | 2.53 | 0.50 ± 0.23 | Plasmid C in communities $B^K$+$B^C$+$B^{CK}$ with 0 $\mu g/ml$ Kanamycin |
| | 1.40 | 0.24 ± 0.28 | Plasmid C in communities $B^K$+$B^C$+$B^{CK}$ with0.5 $\mu g/ml$ Kanamycin |
| | −3.85 | 0.83 ± 0.11 | Plasmid C in communities $B^K$+$B^C$+$B^{CK}$ with 0.5 $\mu g/ml$ Chloramphenicol |

| | | | |
|---|---|---|---|
| | $-0.12$ | $0.84 \pm 0.05$ | Plasmid C in communities $B^{K}$+$B^{C}$+$B^{CK}$ with 2 $\mu g/ml$ Chloramphenicol |
| | $-20.49$ | $0.22 \pm 0.00$ | Plasmid R6K in 3-species (B, R, Y) 3-plasmid (R6K+R388+RP4) communities |
| | $-303.03$ | $0.001 \pm 0.001$ | Plasmid R388 in 3-species (B, R, Y) 3-plasmid (R6K+R388+RP4) communities |
| | $-833.33$ | $0.26 \pm 0.00$ | Plasmid RP4 in 3-species (B, R, Y) 3-plasmid (R6K+R388+RP4) communities |
| Non-transmissible plasmids | $+0$ | $0.04 \pm 0.04$ | Plasmid K– in strain B with 0 $\mu g/ml$ Kanamycin |
| | $-0$ | $0.60 \pm 0.23$ | Plasmid K– in strain B with 0.5 $\mu g/ml$ Kanamycin |
| | $-0$ | $1.00 \pm 0.00$ | Plasmid K– in strain B with 2 $\mu g/ml$ Kanamycin |
| | $+0$ | $0.08 \pm 0.07$ | Plasmid K– in communities $B^{K-}$+$R^{K-}$ with 0 $\mu g/ml$ Kanamycin |
| | $-0$ | $0.08 \pm 0.07$ | Plasmid K– in communities $B^{K-}$+$R^{K-}$ with0.5 $\mu g/ml$ Kanamycin |
| | $-0$ | $1.00 \pm 0.01$ | Plasmid K– in communities $B^{K-}$+$R^{K-}$ with 2 $\mu g/ml$ Kanamycin |

*The plasmid abundance data were extracted from Figures 2, 3, and 4 of this referred paper. Data of 'Simple communities without linoleic acid and phenothiazine' were from Fig. 2c, Fig. 2d, and Fig. 3b. Data of 'Simple communities with linoleic acid and phenothiazine' were from Fig. 4c, Fig. 4d, and Fig. 4e. Data of 'Simple communities with antibiotic treatment' were from Fig. 2c and Fig. 3b. Data of 'Complex communities' were from Fig. 3a, Fig. 3c and Fig. 3d. Data of 'Non-transmissible plasmids' were from Fig. 2b and Fig. 3a.

##### 3.3 Compensatory mutations improve general permissiveness to antibiotic resistance plasmids

In this work, Loftie-Eaton W *et al*. evolved *Pseudomonas sp.H2*, which carries multidrug resistance plasmid RP4, and determined how the capability of plasmid persistence changed with generations. The unstable plasmid-host pair was found to be stabilized due to host adaptation. The ancestral plasmid caused growth burden on the ancestral host, and the plasmid could not be maintained in this ancestral pair. However, after 600 generations, the evolved plasmids became beneficial to the evolved host; this is called plasmid addiction and enabled the plasmid persistence in the evolved pair.

To verify that the change of plasmid persistence during the evolution was governed by a decrease in the $\omega$ value, we calculated the plasmid persistence potential of the ancestral pair (G0) as well as the 600-generation pair (G600), based on the parameter estimations provided by the authors (Table S9). The $\delta$ factor quantifies the difference between the growth rates of plasmid-free cells ($\mu_0$) and plasmid-carrying cells ($\mu_1$) by $\delta = log_2(\mu_1/\mu_0)$. Therefore, the $\lambda$ value in our model can be calculated as $\lambda = 2^{-\delta} - 1$. We estimated the dilution rate to be $0.0375\ h^{-1}$, and normalized the conjugation efficiency with $10^9$. The death rate is much smaller than the growth rate, thus the term $\frac{\mu}{\mu-d}$ was approximated as 1 in our calculation. The $\omega$ values and the corresponding relative abundance of plasmid RP4 are summarized in Table S10.

**Table S9** Original data of plasmid RP4 relative abundance from 'Compensatory mutations improve general permissiveness to antibiotic resistance plasmids'

| | | | Replicate 1 | Replicate 2 | Replicate 3 |
|---|---|---|---|---|---|
| Ancestral pair (G0) | Population A | | 3/52 | 8/52 | 6/52 |
| | Population B | | 4/52 | 4/52 | 0/52 |
| | Population C | | 1/52 | 1/52 | 0/52 |
| 600th generation (G600) | Population A | Clone A1 | 52/52 | 52/52 | 52/52 |
| | | Clone A2 | 49/52 | 52/52 | 52/52 |
| | | Clone A3 | 52/52 | 51/52 | 51/52 |
| | Population B | Clone B1 | 52/52 | 52/52 | 52/52 |
| | | Clone B2 | 51/52 | 50/52 | 52/52 |
| | | Clone B3 | 52/52 | 52/52 | 52/52 |
| | Population C | Clone C1 | 50/52 | 50/52 | 50/52 |
| | | Clone C2 | 44/52 | 49/52 | 47/52 |
| | | Clone C3 | 49/52 | 43/52 | 40/52 |

*The plasmid abundance data, which were visualized in Fig. 1 of this referred paper, were obtained from DRYAD (https://datadryad.org/bitstream/handle/10255/dryad.147859/ResultsSection2.1.csv?sequence=1)

**Table S10** The parameter estimations, plasmid RP4 persistence potential and the corresponding plasmid relative abundance in 'Compensatory mutations improve general permissiveness to antibiotic resistance plasmids'

| | | | $\delta$ | $\kappa$ | $\eta$ | $\omega$ | $f$ |
|---|---|---|---|---|---|---|---|
| | | | Plasmid effect on fitness | Plasmid segregation loss rate | Plasmid conjugation efficiency | Plasmid persistence potential | Plasmid relative abundance |
| Ancestral pair (G0) | Population A | | $-6.59\ \%$ | $10^{-7.21}$ | $10^{-12.85}$ | 0.084 | $0.11 \pm 0.05$ |
| | Population B | | $-7.75\ \%$ | $10^{-6.51}$ | $10^{-12.72}$ | 0.097 | $0.05 \pm 0.04$ |
| | Population C | | $-7.91\ \%$ | $10^{-6.31}$ | $10^{-12.80}$ | 0.079 | $0.01 \pm 0.01$ |
| 600th generation (G600) | Population A | Clone A1 | 2.48 % | $10^{-3.32}$ | $10^{-13.54}$ | $-0.17$ | $1 \pm 0.00$ |
| | | Clone A2 | 2.79 % | $10^{-3.38}$ | $10^{-12.87}$ | $-0.43$ | $0.98 \pm 0.03$ |
| | | Clone A3 | 2.56 % | $10^{-3.89}$ | $10^{-12.92}$ | $-0.22$ | $0.99 \pm 0.01$ |
| | Population B | Clone B1 | 11.09 % | $10^{-3.70}$ | $10^{-12.14}$ | $-0.26$ | $1 \pm 0.00$ |

| | | | | | | | |
|---|---|---|---|---|---|---|---|
| | | Clone B2 | 6.43 % | $10^{-3.61}$ | $10^{-12.95}$ | $-0.077$ | $0.98 \pm 0.02$ |
| | | Clone B3 | 8.37 % | $10^{-4.08}$ | $10^{-12.04}$ | $-0.42$ | $1 \pm 0.00$ |
| | Population C | Clone C1 | 10.16 % | $10^{-2.87}$ | $10^{-12.69}$ | $-0.15$ | $0.96 \pm 0.00$ |
| | | Clone C2 | 12.40 % | $10^{-2.65}$ | $10^{-13.41}$ | $-0.035$ | $0.90 \pm 0.05$ |
| | | Clone C3 | 8.45 % | $10^{-2.84}$ | $10^{-12.92}$ | $-0.15$ | $0.85 \pm 0.09$ |

*The plasmid effects on fitness ($\delta$) were extracted from Fig. 3a of this referred paper. The plasmid segregation loss rates ($\kappa$) were extracted from Fig. 3b of this referred paper. The plasmid conjugation efficiencies ($\eta$) were extracted from Fig. S2 of this referred paper. We changed the letters of each parameter to fit the denotations of our model.

##### 3.4 Positive selection inhibits gene mobilization and transfer in soil bacterial communities

Hall JP *et al.* conducted a similar coculture experiment using *Pseudomonas fluorescens*, *Pseudomonas putica*, and plasmid pQBR57 as described in 3.1. We focused only on the data after transfer 29. We borrowed the $\omega$ values at different transfers from Table S6 and calculated the plasmid relative abundance from their original data (Table S11). Notably, in Fig. 2 of their main text, they also showed the results of short-term dynamics (transfer 0 to 5) of plasmid pQBR57 in *Pseudomonas fluorescens* cultured alone. However, we did not consider these results because at the end of the experiments (transfer 5), the population was far from the steady state.

**Table S11** The plasmid pQBR57 persistence potential and the corresponding relative abundance in 'Positive selection inhibits gene mobilization and transfer in soil bacterial communities'

| | Plasmid persistence potential | Plasmid abundance |
|---|---|---|
| Transfer 29 | $1.39 \pm 0.08$ | 0.96875 |
| Transfer 35 | $1.41 \pm 0.09$ | 0.96875 |
| Transfer 41 | $1.39 \pm 0.05$ | 0.90625 |
| Transfer 47 | $1.42 \pm 0.06$ | 0.93750 |
| Transfer 53 | $1.34 \pm 0.10$ | 0.81250 |
| Transfer 59 | $1.38 \pm 0.06$ | 0.96875 |
| Transfer 65 | $1.13 \pm 0.13$ | 0.34375 |

*The plasmid persistence potentials were calculated from kinetic parameters given in Table S5. The plasmid abundance data, which were visualized in Fig. 3b of this referred paper, was obtained from DRYAD (https://datadryad.org/bitstream/handle/10255/dryad.148630/S4_Tn5042_PF_Data.csv?sequence=1).

##### 3.5 Parallel compensatory evolution stabilizes plasmids across the parasitism-mutualism continuum

Harrison E. *et al.*, investigated the compensatory evolution across the parasitism-mutualism continuum. They established 36 populations of bacterium *Pseudomonas fluorescens* SBW25 carrying plasmid QBR103, which encodes mercury resistance, and propagated the populations by serial transfer under six mercury concentrations ($0, 8, 16, 24, 32,$ and $40$ $\mu$M $HgCl_2$). They analyzed the fitness of the ancestral and evolved generations in each environment as well as the conjugation rates. These data, combined with the plasmid abundance at the end of the transfers in each environment, are summarized in Table S12.

The fitness value $\alpha$ of the evolved host-plasmid pair relative to the plasmid-free cells can be calculated by $\alpha = \alpha_A \alpha_E$, where $\alpha_A$ is the fitness of the ancestral pair relative to the plasmid-free cells and $\alpha_E$ is the fitness of the evolved pair relative to the ancestors. The values of $\alpha_E$ are summarized in Table S12. However, the author did not directly provide the measured values of $\alpha_A$ under the six $HgCl_2$ concentrations. Instead, they provided a linear correlation between $\alpha_A$ and $HgCl_2$ concentrations with the unit of $\mu$M: $\alpha_A \approx 0.0371 \times [HgCl_2] + 0.3019$ (Figure S2 in the Supplementary Information of their article). In this way, we were able to calculate the $\alpha$ values in the six cases. The $\lambda$ values in our model can then be calculated by $\lambda = \alpha^{-1} - 1$. We estimated the dilution rate $D$ of the transfers to be 0.0125. To become a dimensionless quantity, the conjugation efficiencies are required to be normalized with the maximum carrying capacity $N_m$. However, the authors did not provide this value in the article. Using similar experimental settings, Hall JP estimated the maximum carrying capacity of *Pseudomonas fluorescens* to be $6.01 \times 10^8$, as is shown in Table S5, and we used this estimation in this analysis. We also used Hall's estimation of segregation rate $\kappa \approx 1 \times 10^{-4} h^{-1}$. The dilution rate is much smaller than the growth rate, thus the plasmid persistence potential can be approximated by $\omega \approx \bar{\eta} / \left( D + \bar{\kappa} - \frac{D}{1+\bar{\lambda}} \right)$. Our estimations of persistence potential and the corresponding plasmid abundance are summarized in Table S12.

**Table S12** Kinetic parameters, persistence potential and plasmid abundance of plasmid pQBR103 in 'Parallel compensatory evolution stabilizes plasmids across the parasitism-mutualism continuum'

| $HgCl_2$ concentration ($\mu$M) | Populations | $\alpha_E$: Evolved fitness relative to ancestors | $ln$(conjugation rate) | Plasmid persistence potential | Plasmid relative abundance |
|---|---|---|---|---|---|
| 0 | A | 1.30 | −14.92 | $-1.05 \times 10^4$ | 0.701 |
| | B | 1.20 | −15.77 | $-3.87 \times 10^3$ | 0.601 |
| | C | 1.51 | −15.80 | $-5.55 \times 10^3$ | 0.403 |
| | D | 0.97 | −15.97 | $-2.33 \times 10^3$ | 0.252 |
| | E | 1.27 | −17.03 | $-1.21 \times 10^3$ | 0.094 |
| | F | 1.44 | −16.90 | $-1.70 \times 10^3$ | 0.003 |
| 8 | A | 1.80 | −17.44 | $1.61 \times 10^4$ | 1 |

| | | | | | |
|---|---|---|---|---|---|
| | B | 1.39 | −17.94 | $-4.02\times10^3$ | 1 |
| | C | 1.22 | −15.30 | $-3.05\times10^4$ | 1 |
| | D | 1.17 | −15.34 | $-2.53\times10^4$ | 1 |
| | E | 1.08 | −15.01 | $-2.71\times10^4$ | 1 |
| | F | 1.38 | −16.63 | $-1.43\times10^4$ | 0.201 |
| 16 | A | 1.09 | −14.07 | $-2.44\times10^6$ | 1 |
| | B | 1.15 | −15.82 | $1.85\times10^5$ | 1 |
| | C | 1.33 | −15.69 | $4.30\times10^4$ | 1 |
| | D | 1.57 | −15.86 | $2.09\times10^4$ | 1 |
| | E | 1.33 | −13.90 | $2.62\times10^5$ | 1 |
| | F | 1.41 | −17.61 | $5.05\times10^3$ | 1 |
| 24 | A | 1.20 | −15.41 | $3.16\times10^4$ | 0.901 |
| | B | 1.38 | −15.37 | $2.54\times10^4$ | 1 |
| | C | 1.22 | −15.49 | $2.77\times10^4$ | 1 |
| | D | 1.33 | −15.97 | $1.47\times10^4$ | 1 |
| | E | 1.19 | −16.60 | $9.71\times10^3$ | 1 |
| | F | 1.46 | −17.75 | $2.17\times10^3$ | 1 |
| 32 | A | 1.35 | −14.58 | $4.40\times10^4$ | 1 |
| | B | 1.40 | −14.68 | $3.84\times10^4$ | 1 |
| | C | 1.23 | −15.92 | $1.27\times10^4$ | 1 |
| | D | 1.11 | −16.49 | $8.22\times10^3$ | 1 |
| | E | 1.29 | −17.62 | $2.20\times10^3$ | 1 |
| | F | 1.39 | −17.93 | $1.50\times10^3$ | 1 |
| 40 | A | 2.07 | −15.07 | $1.86\times10^4$ | 1 |
| | B | 1.61 | −15.45 | $1.42\times10^4$ | 1 |
| | C | 1.21 | −16.51 | $5.96\times10^3$ | 1 |
| | D | 1.37 | −16.82 | $3.99\times10^3$ | 1 |
| | E | 1.22 | −16.77 | $4.54\times10^3$ | 1 |
| | F | 1.68 | −17.48 | $1.83\times10^3$ | 1 |

*The $\alpha_E$ values were extracted from Fig. 1A of this referred article. We changed the letters of the parameter to fit the denotations of in our model. The conjugation rates were extracted from Fig. 2B of this referred article. The fitness of the ancestral pair relative to the plasmid-free cells were extracted from Fig. S1 in the Supplementary Information, and the data of plasmid abundance were extracted from Fig. S2 of this referred article.

**3.6 Variable plasmid fitness effects and mobile genetic element dynamics across *Pseudomonas* species**

Kottara A *et al*. tracked the dynamics of a large conjugative plasmid, pQBR103 (mercury-resistant), across five diverse *Pseudomonas* species (*P. fluorescens*, *P. savastanoi*, *P. stutzeri*, *P. aeruginosa*, and *P. putida*) in environments with and without mercury selection. They measured the fitness of the *Pseudomonas* species carrying the plasmid as well as the long-term frequencies of $Hg^R$ phenotypes with or without 50 $\mu$M mercury. The pQBR103 conjugation efficiency of *P. fluorescens* can be obtained as $e^{-14.66}$ from Harrison E *et al*., 2015. The conjugation efficiencies of the other four species were lacking. Therefore, we only considered the data of *P. fluorescens*.

The $\lambda$ value in our model was calculated from the selection rate $\gamma$ through $\lambda = 2^{-\gamma} - 1$. We normalized the conjugation efficiency of *P. fluorescens* with respect to the carrying capacity $6.01 \times 10^8$ (Table S5). We estimated the plasmid segregation rate $\kappa$ to be $1 \times 10^{-4}\ h^{-1}$ and the cellular death rate to be $0.009\ h^{-1}$ (Table S5). When calculating the persistence potential $\omega = \frac{\bar{\eta}}{\frac{\bar{\mu}}{\bar{\mu}-\sigma d}\left(d+\bar{\kappa}-\frac{d}{1+\bar{\lambda}}\right)}$, we approximated the term $\frac{\bar{\mu}}{\bar{\mu}-\sigma d}$ to be 1 since the growth rate was much greater than the death rate. Therefore, we obtained the plasmid persistence potential as $\bar{\eta}/\left(d+\bar{\kappa}-\frac{d}{1+\bar{\lambda}}\right)$. The $\omega$ values and the corresponding plasmid relative abundance data are summarized in Table S13.

**Table S13** Kinetic parameters, persistence potential and relative abundance of plasmid pQBR103 in 'Variable plasmid fitness effects and mobile genetic element dynamics across *Pseudomonas* species'

| | Description | Value |
|---|---|---|
| $\gamma_0$ | The selection rate of P. Fluorescens carrying plasmid pQBR103 without mercury selection | $-0.425$ |
| $\gamma_{50}$ | The selection rate of P. Fluorescens carrying plasmid pQBR103 with 50 $\mu$M mercury selection | 2.053 |
| $\omega_0$ | Persistence potential of plasmid pQBR103 without mercury selection | $1.08 \times 10^5$ |
| $\omega_{50}$ | Persistence potential of plasmid pQBR103 with 50 $\mu$M mercury selection | $-9.14 \times 10^3$ |
| $f_{60}^0$ | Plasmid pQBR103 abundance without selection at transfer 60 | $0.69 \pm 0.17$ |
| $f_{48}^0$ | Plasmid pQBR103 abundance without selection at transfer 48 | $0.75 \pm 0.06$ |
| $f_{36}^0$ | Plasmid pQBR103 abundance without selection at transfer 36 | $0.83 \pm 0.09$ |
| $f_{60}^{50}$ | Plasmid pQBR103 abundance with 50 $\mu$M mercury selection at transfer 60 | $0.75 \pm 0.03$ |
| $f_{48}^{50}$ | Plasmid pQBR103 abundance with 50 $\mu$M mercury selection at transfer 48 | $0.81 \pm 0.08$ |
| $f_{36}^{50}$ | Plasmid pQBR103 abundance with 50 $\mu$M mercury selection at transfer 36 | $0.82 \pm 0.04$ |

*The $\gamma$ values were extracted from Fig. 1 of this referred article. The data of plasmid abundance were extracted from Fig. 2 of this referred article.

##### 3.7 Amelioration of the cost of conjugative plasmid carriage in *Escherichia coli* K12

Dahlberg C *et al.* studied how the cost of conjugative plasmids carrying drug resistance evolved in batch cultures in the absence of antibiotic selection. They studied two plasmids, R1 and RP4, both of which carried multiple drug resistance genes and imposed an initial fitness cost on the host *Escherichia coli*. To determine whether the fitness cost could be reduced, they subjected the plasmid-carrying bacteria to 1100 generations of evolution. They then measured the changes in fitness costs as well as the conjugation efficiencies.

The plasmid-abundance data of RP4 were not presented in the paper, so we only considered the plasmid R1. The fitness costs, conjugation rates, and plasmid abundance of three evolved R1 populations are shown in Table S14. The $\lambda$ factors can be obtained from the fitness cost $\alpha$ through $\lambda = \alpha - 1$. We normalized the conjugation efficiency using the maximum carrying capacity $N_m =$

$8 \times 10^9$. We estimated the segregation rate to be $0.001$, and the dilution rate $D$ to be $0.025$. We also assumed the growth rates to be much larger than the dilution rate, so that the factor $\frac{\mu}{\mu - D}$ was approximated as 1. The values of plasmid persistence potential $\omega$ are also summarized in Table S14.

**Table S14** Kinetic parameters, persistence potential and relative abundance of plasmid R1 in 'Amelioration of the cost of conjugative plasmid carriage in *Eschericha coli* K12'

| | Population 1 | Population 2 | Population 3 |
|---|---|---|---|
| Fitness cost $\alpha$ | $1.286$ | $1.303$ | $1.309$ |
| Conjugation efficiency $\eta$ | $2.25 \times 10^{-12}$ | $1.14 \times 10^{-12}$ | $9.14 \times 10^{-12}$ |
| Plasmid fraction | $105/106$ | $97/106$ | $66/88$ |
| Plasmid persistence potential | $2.74$ | $1.34$ | $10.59$ |

*The $\alpha$ values were extracted from Table 2 of this referred article. Population 1, 2, 3 correspond to strain CD111, CD112, and CD113, respectively. The conjugation rates were obtained from Table 4, and the plasmid fractions were obtained from Table 3.

##### 3.8 The IncI1 plasmid carrying the $bla_{CTX\text{-}M\text{-}1}$ gene persists in in vitro culture of an *Escherichia coli* strain from broilers

Fischer EA *et al.* studied the conjugation dynamics of the lncl1 plasmid carrying *$bla_{CTX\text{-}M\text{-}1}$* gene in a batch culture. They quantified the population dynamics of three *E. coli* populations (donors, recipients, and transconjugants) in two mixed-culture experiments. In the first experiment, donor cells were mixed with recipient cells and the mixture was incubated for 24 hours without dilution. To determine the cell densities of donors, recipients, and transconjugants, the samples were taken out for colony counts at 0, 3, 6, 16, 19, and 24 h after the start of experiments. In the second experiment, transconjugants and recipients were mixed, and the culture were passaged every 24 hours or 48 hours with a dilution ratio of 1:10,000. The cultures were passaged over a period of three months. The cell densities of recipients and transconjugants were measured through plating and colony counts. The plasmid abundance data and the estimations of kinetic parameters are summarized in Table S15.

In the first experiment, there was no dilution ($D = 0$). Therefore, the plasmid persistence potential $\omega$ can be obtained as $\omega = \frac{\bar{\eta}}{\kappa}$. The plasmid loss rate was provided as $0.0008 \sim 0.0036$. We used the median value $0.0022$ as the estimation of $\kappa$. We calculated $\bar{\eta}$ through $\bar{\eta} = \frac{(\eta_D N_m)N_D + (\eta_T N_m)N_T}{N_D + N_R + N_T}$, where $N_D$, $N_R$, and $N_T$ are the cell densities of donors, recipients, and transconjugants, respectively. $\eta_D$ and $\eta_T$ are the conjugation rates of donors and transconjugants, respectively. Here we normalized the conjugation rates with respect to the maximum carrying capacity $N_m$.

In the second experiment, the conjugation efficiency became $\eta = \eta_T N_m$. For the dilution period of 24 hours, we estimated the dilution rate $D = 0.05$, and for the dilution period of 48 hours, we estimated $D = 0.025$. The $\lambda$ factor could be obtained through $\lambda = \frac{\mu_R}{\mu_D} - 1$, where $\mu_R$ and $\mu_D$ are the growth rates of donors and recipients, respectively. The $\omega$ values in both cases could then be calculated through $\omega = \frac{\eta}{\frac{\mu_R}{\mu_R - D}\left(D + \kappa - \frac{D}{1+\lambda}\right)}$. The $\omega$ values and the corresponding plasmid abundances are summarized in Table S15.

**Table S15** Kinetic parameters, persistence potential and relative abundance of IncI1 plasmid in 'The IncI1 plasmid carrying the *bla*CTX-M-1 gene persists in *in vitro* culture of an *Escherichia coli* strain from broilers'

<table>
<tr><th colspan="3">Description</th><th>Value</th></tr>
<tr><td colspan="3">Growth rate of recipient cells $\mu_R$</td><td>$2.04\ h^{-1}$</td></tr>
<tr><td colspan="3">Growth rate of donor cells $\mu_D$</td><td>$2.09\ h^{-1}$</td></tr>
<tr><td colspan="3">Growth rate of transconjugant cells $\mu_T$</td><td>$2.09\ h^{-1}$</td></tr>
<tr><td colspan="3">Maximum carrying capacity of the mixture $N_m$</td><td>$9.33 \times 10^8\ cfu/ml$</td></tr>
<tr><td colspan="3">Plasmid loss rate $\kappa$</td><td>$0.008 \sim 0.0036\ h^{-1}$</td></tr>
<tr><td colspan="3">Conjugation efficiency of donor cells $\eta_D$</td><td>$2.4 \times 10^{-14}\ cfu^{-1}h^{-1}ml$</td></tr>
<tr><td colspan="3">Conjugation efficiency of transconjugant cells $\eta_T$</td><td>$4.4 \times 10^{-10}\ cfu^{-1}h^{-1}$</td></tr>
<tr><td colspan="2" rowspan="5">Short-term experiments without dilution</td><td>Population density of donors $N_D$</td><td>$4.02 \times 10^8$</td></tr>
<tr><td>Population density of recipients $N_R$</td><td>$6.19 \times 10^8$</td></tr>
<tr><td>Population density of transconjugants $N_T$</td><td>$6.19 \times 10^7$</td></tr>
<tr><td>Plasmid persistence potential $\omega$</td><td>117.34</td></tr>
<tr><td>Plasmid abundance $f$</td><td>0.43</td></tr>
<tr><td rowspan="4">Long-term experiments with dilution</td><td rowspan="2">Dilution every 24 hours</td><td>Plasmid persistence potential $\omega$</td><td>410.93</td></tr>
<tr><td>Plasmid abundance $f$</td><td>$1.03 \pm 0.28$</td></tr>
<tr><td rowspan="2">Dilution every 48 hours</td><td>Plasmid persistence potential $\omega$</td><td>255.47</td></tr>
<tr><td>Plasmid abundance $f$</td><td>$0.94 \pm 0.28$</td></tr>
</table>

*The values of $\mu_R$, $\mu_D$, $\mu_T$ were obtained from Table 1 of this referred paper. The value of $N_m$ was obtained from Table 2. The value of $\kappa$ was obtained from the main text. The values of $\eta_D$ and $\eta_T$ were obtained from Table 3. The population density data of short-term experiments were extracted from Fig. 2 of this referred paper. The plasmid abundance data of long-term experiments were extracted from Fig. 3. We changed the letters of each parameters to fit the denotations of our model.

##### 3.9 Survival and evolution of a large multidrug resistance plasmid in new clinical bacterial hosts

Porse A *et al.* investigated the survival and evolution of a large multidrug resistance plasmid, pKp33, in three clinical bacterial hosts: Ec37, Ec38 and Kp08. They evolved the host-plasmid pairs for ~280 generations and then measured the plasmid stability of both naïve and evolved host-plasmid pairs. During the measurements, the plasmid dynamics in evolved host-plasmid pairs did not reach steady states, thus we only considered the dynamics of naïve plasmid-host pairs. The values of kinetic parameters are summarized in Table S16.

During the experiments, they adopted a daily dilution ratio of 1:150. Therefore, we estimated the dilution rate $D$ to be $0.0272$. We approximated the plasmid persistence potential as $\omega = \eta / \left(D + \kappa - \frac{D}{1+\lambda}\right)$, since the dilution rate is much smaller than the growth rate. $\lambda$ values can be obtained from $\rho$ through $\lambda = \frac{1}{1-\rho} - 1$. We also normalized the conjugation efficiency with the maximum carrying capacity $N_m = 1 \times 10^9$ . The plasmid persistence potentials and the corresponding plasmid abundance data are summarized in Table S16.

**Table S16** Kinetic parameters, plasmid persistence potential and plasmid relative abundance data in 'Survival and evolution of a large multidrug resistance plasmid in new clinical bacterial hosts'

| | Ec37/pKP33 | Ec38/pKP33 | Kp08/pKP33 |
|---|---|---|---|
| Conjugation rate $\eta$ | $5.03 \times 10^{-14}$ | $4.21 \times 10^{-13}$ | $7.17 \times 10^{-14}$ |
| Segregation rate $\kappa$ | 0.0147 | 0.0008 | 0.0060 |
| Plasmid cost $\rho$ | 8.3% | 14.0% | 4.6% |
| Plasmid persistence potential $\omega$ | 0.003 | 0.091 | 0.99 |
| Plasmid relative abundance | 0 | 0 | 0 |

*The values of conjugation rates were obtained from Supplementary Table S4 of this referred paper. The values of segregation rates and plasmid costs were obtained from Supplementary Table S5 of this referred paper. The plasmid abundance data of were extracted from Fig. 4. We changed the letters of each parameter to fit the denotations of our model.

##### 3.10 Resource-limited growth, competition, and predation: a model and experimental studies with bacteria and bacteriophage

Levin BR *et al.* presented models of phage-bacteria population growth. Based on assumptions of predator-prey interactions, they predicted the conditions of stable coexistence. They then compared the behavior predicted by the models with that of the experimental chemostat populations of *E. coli* and its virulent virus T2. First, they investigated the population with the bacteriophage T2 and a T2-sensitive strain of *E. coli* in glucose-limited conditions with two different dilution rates. In both cases, there were equilibria with the phage and bacterial populations persisting. They then investigated the population with T2, one T2-sensitive strain, and one T2-resistant strain, and they observed an equilibrium with three populations coexisting.

The persistence potential of the phage was calculated as $\omega = \frac{\eta}{\frac{\mu}{\mu - D}\cdot(D+\kappa)}$. The rate of exponential growth is represented by $\mu$. $D$ is the dilution rate. Unlike in the scenarios of plasmids and transposons, $\kappa$ here represents the host mutation rates to phage resistance, which was estimated to be extremely small compared with $D$[16]. Here $\eta$ represents the infection rates of phages. We calculated $\eta = \frac{\delta\beta}{\theta} N_m$, where $\delta$ stands for the phage absorption rate, $\beta$ stands for the phage burst size, $\theta$ stands for the latent time of infection, and $N_m$ stands for the maximum carrying capacity of the bacteria. $N_{cell}$ represents the cell density, and $N_{phage}$ represents the phage density. We defined the phage abundance as $\frac{N_{phage}}{N_{phage}+\beta N_{cell}}$. The parameters and phage abundance data are summarized in Table S17.

**Table S17** Kinetic parameters, phage persistence potential and phage relative abundance data in 'Resource-Limited Growth, Competition, and Predation: A Model and Experimental Studies with Bacteria and Bacteriophage'

| description | | | value |
|---|---|---|---|
| rate of exponential growth ($\mu$) | | | $0.738\ h^{-1}$ |
| phage absorption rate ($\delta$) | | | $6.24 \times 10^{-8}\ ml \cdot h^{-1}$ |
| phage burst size ($\beta$) | | | 98 |
| latent time of infection ($\theta$) | | | $0.5\ h$ |
| host mutation rate to phage resistance | | | $8 \times 10^{-8}\ h^{-1}$ |
| one-prey experiments | $D = 0.130\ h^{-1}$ | cell density | $2.88 \times 10^{4}\ ml^{-1}$ |
| | | phage density | $2.73 \times 10^{6}\ ml^{-1}$ |
| | | phage persistence potential | 2.24 |
| | | phage abundance | 0.49 |
| | $D = 0.217\ h^{-1}$ | cell density | $4.95 \times 10^{3}\ ml^{-1}$ |
| | | phage density | $1.00 \times 10^{6}\ ml^{-1}$ |
| | | phage persistence potential | 1.15 |
| | | phage abundance | 0.67 |
| two-prey experiment | dilution rate ($D$) | | $0.269\ h^{-1}$ |
| | sensitive cell density | | $5.99 \times 10^{6}\ ml^{-1}$ |
| | resistant cell density | | $2.30 \times 10^{8}\ ml^{-1}$ |
| | phage density | | $6.49 \times 10^{7}\ ml^{-1}$ |
| | phage persistence potential | | 0.98 |
| | phage abundance | | 0.0028 |

* The parameters were obtained from the main text of this referred paper. The abundance data of bacteriophage T2 were abstracted from Fig. 5 and Fig. 8 of this referred paper.

##### 3.11 Constraints on the coevolution of bacteria and virulent phage: a model, some experiments, and predictions for natural communities

Lenski RE *et al.* expanded Levin's work (as described in section 3.10) and incorporated mutational events into the population dynamics. Here we use their data of the experiments with *E. coli* and phage T4 in chemostat for our analysis. Our calculation process was described in 3.10. The parameters and phage abundance data are summarized in Table S18.

**Table S18** Kinetic parameters, phage persistence potential and phage relative abundance data in 'Constraints on the Coevolution of Bacteria and Virulent Phage: A Model, Some Experiments, and Predictions for Natural Communities'

| description | value |
|---|---|
| rate of exponential growth ($\mu$) | $0.7\ h^{-1}$ |
| phage absorption rate ($\delta$) | $3 \times 10^{-7}\ ml \cdot h^{-1}$ |
| phage burst size ($\beta$) | 80 |
| latent time of infection ($\theta$) | $0.6\ h$ |
| host mutation rate to phage resistance | $8 \times 10^{-8}\ h^{-1}$ |

| | |
|---|---|
| cell density | $1.72 \times 10^{8}\ ml^{-1}$ |
| phage density | $5.60 \times 10^{3}\ ml^{-1}$ |
| phage persistence potential | $1.46 \times 10^{-5}$ |
| phage abundance | $4.07 \times 10^{-7}$ |

* The parameters were obtained from the Table 1 of this referred paper. The abundance data of bacteriophage T4 were abstracted from Fig. 1 of this referred paper.

##### 3.12 Effect of resource enrichment on a chemostat community of bacteria and bacteriophage

Bohannan BJ *et al*. investigated how resource enrichment influences the population equilibrium of *E. coli* and bacteriophage T4. They supplied the community in chemostat with different concentrations of glucose and tracked the long-term population dynamics. We used the parameter values provided by Lenski *et al.* 1985 (Table S18). The data of population densities and phage abundances were summarized in Table S19.

**Table S19** phage persistence potential and phage relative abundance data in 'Effect of resource enrichment on a chemostat community of bacteria and bacteriophage'

| description | | value |
|---|---|---|
| 0.1 mg/L glucose | dilution rate ($D$) | $0.3\ h^{-1}$ |
| | total cell density | $7.42 \times 10^{4}\ ml^{-1}$ |
| | resistant cell density | $3.42 \times 10^{3}\ ml^{-1}$ |
| | phage density | $4.01 \times 10^{4}\ ml^{-1}$ |
| | phage persistence potential | 0.16 |
| | phage abundance | 0.0067 |
| 0.5 mg/L glucose | dilution rate ($D$) | $0.3\ h^{-1}$ |
| | total cell density | $6.35 \times 10^{5}\ ml^{-1}$ |
| | resistant cell density | $2.43 \times 10^{5}\ ml^{-1}$ |
| | phage density | $2.12 \times 10^{4}\ ml^{-1}$ |
| | phage persistence potential | 0.73 |
| | phage abundance | $4.16 \times 10^{-4}$ |

* The parameters were borrowed from Lenski *et al.* 1985[33]. The abundance data of bacteriophage T4 were abstracted from Fig. 3 of this referred paper.

##### 3.13 The evolution of transposable elements: conditions for establishment in bacterial populations

Condit R studied the conditions for the establishment of transposons in *E. coli* populations using the conjugative plasmid R100 and the transposons Tn3 and Tn5. A plasmid-borne transposon was introduced into the *E. coli* population carrying the same plasmid without the transposon. The proportion of the transposon was tracked long-term. Four groups of experiments were done, but for two of them the abundance data at the end of the long-term experiments were not representative of the steady state. Thus, we only used the data from two experiments, represented in Figs. 3 and 4.

The transposon abundance data as well as kinetic parameters are summarized in Table S20. As estimated by the author, the mean transposition rates for Tn5 and Tn3 were $7.1 \times 10^{-7}\ hr^{-1}$ and $6.3 \times 10^{-6}\ hr^{-1}$, respectively, while the conjugation efficiencies were no higher than

$2.4 \times 10^{-11}$. Therefore, conjugation efficiencies were negligible compared with the transposition rates. We applied the quasi-equilibrium assumption for the transposition process. The author assumed that the plasmid-to-chromosome transposition rate $\tau_C$ was the same as the chromosome-to-plasmid rate $\tau_P$. Thus, we estimated the value of the equilibrium constant $K = \tau_P/\tau_C$ as $K = 1$. The transposon transfer rate $\eta$ is estimated from $\eta_0$ through $\eta = \eta_0 K(1 + K)$. The growth burden $\lambda$ is calculated from the relative fitness $W$ through $\lambda = W^{-1} - 1$. For the experiments of daily transfers in minimal medium, the dilution ratio was 1:100; therefore, we estimated the dilution rate $D = \frac{ln10^2}{24\ hr} = 0.192\ hr^{-1}$. The transposon loss rate was estimated to be extremely small compared with the dilution rate. Thus, we approximate the persistence potential as $\omega = \frac{\eta N}{\frac{\mu}{\mu - D}(D - \frac{D}{1+\lambda})}$. For the chemostat experiments, the dilution rate was 0.13 $hr^{-1}$, but the maximum growth rate was not provided. We assumed that the maximum growth rate was much higher than 0.13 $hr^{-1}$ based on the growth rates in minimal medium and Luria Broth. Thus, we approximate the persistence potential in chemostat as $\omega = \frac{\eta N}{D - \frac{D}{1+\lambda}}$. The persistence potentials are also summarized in Table S20.

**Table S20** The transposon abundance data and kinetic parameters in 'The evolution of transposable elements: conditions for establishment in bacterial populations'

| Parameters | | Values |
|---|---|---|
| Minimal Medium | Growth rate ($\psi$) | 0.72 $hr^{-1}$ |
| | Maximum cell density ($N$) | $5.1 \times 10^8$ cell $ml^{-1}$ |
| | Relative fitness ($W$) | 0.94 |
| | Dilution rate ($D$) | 0.192 $hr^{-1}$ |
| | Conjugation rate ($\eta_0$) | $2.4 \times 10^{-11}$ $cell^{-1}$ ml $hr^{-1}$ |
| | Persistence potential of Tn3 | 0.39 |
| | Relative abundance of Tn3 | $2.8 \times 10^{-6}$ |
| Chemostat | Maximum cell density ($N$) | $3.0 \times 10^9$ cell $ml^{-1}$ |
| | Relative fitness ($W$) | 0.85 |
| | Dilution rate ($D$) | 0.192 $hr^{-1}$ |
| | Conjugation rate of R3 ($\eta_{03}$) | $2.4 \times 10^{-13}$ $cell^{-1}$ ml $hr^{-1}$ |
| | Conjugation rate of R5 ($\eta_{05}$) | $7.0 \times 10^{-17}$ $cell^{-1}$ ml $hr^{-1}$ |
| | Persistence potential of Tn3 | 0.043 |
| | Relative abundance of Tn3 | $2.0 \times 10^{-6}$ |
| | Persistence potential of Tn5 | $1.9 \times 10^{-5}$ |
| | Relative abundance of Tn5 | 0.0012 |

* The parameters were obtained from Tables 1 and 2 of this referred paper. The abundance data of the transposons were abstracted from Figs. 3 and 4 of this referred paper.

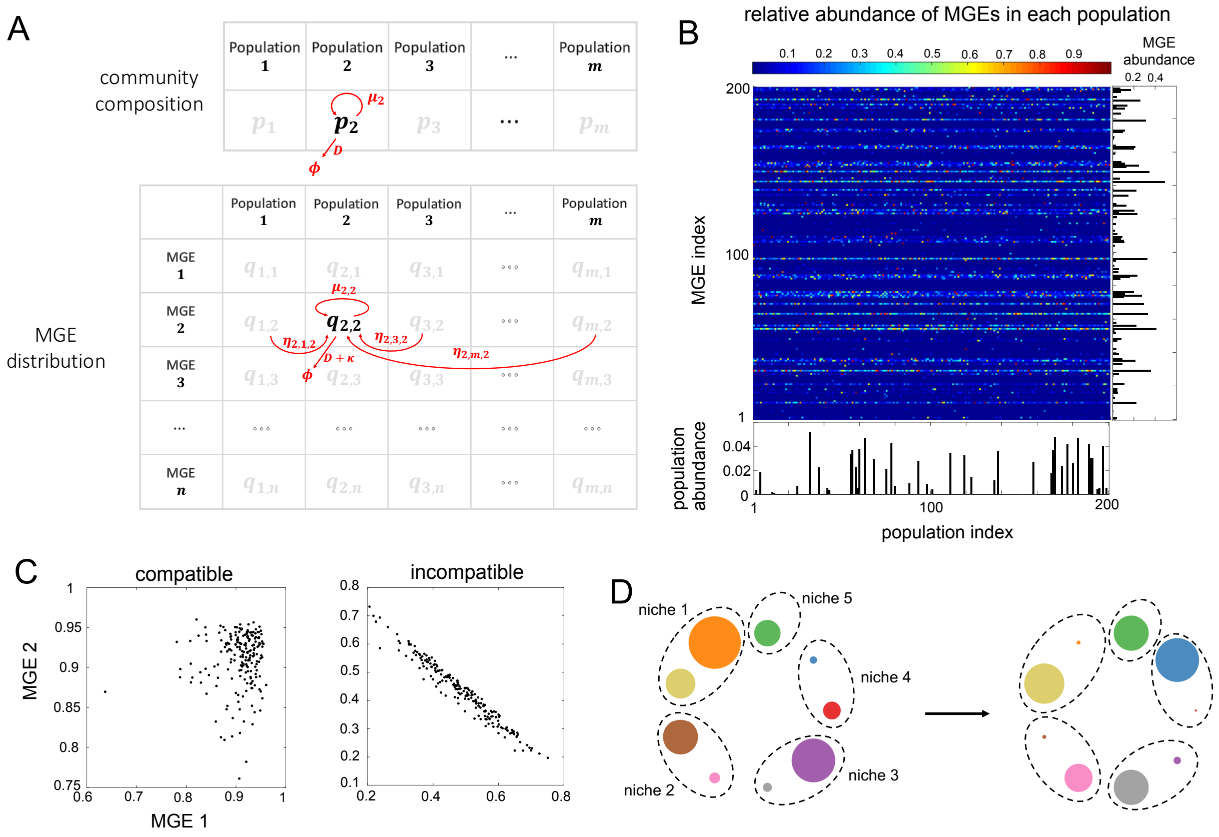


**Fig. S1. MGE-centric framework (MCF) and its application in predicting microbial community dynamics.**

(**A**) Dynamic processes in MCF. The variables in MCF include the species abundance ($s$) and MGE abundance in each species ($p$). Four processes are described in this framework: cell division ($\mu$), dilution ($D$), MGE horizontal transfer ($\eta$), and MGE loss ($\kappa$).

(**B**) MCF is capable of predicting the steady-state composition, MGE abundance, and distribution patterns of huge communities. A community with 200 species and 200 MGEs is shown as an example here. All parameters are randomized. The relative abundance of MGE $j$ in species $i$ was calculated as the fraction of species $i$ cells that contains MGE $j$ relative to the total number of species $i$ cells.

(**C**) MCF can be applied to incompatible plasmids. A community of 200 species and 2 plasmids, compatible or incompatible with each other, are shown as an example. In each species, the relative abundance of the plasmids, calculated as the fraction of the plasmid-carrying cells, are calculated and plotted. The simulation was carried out with the randomized parameters within the following ranges: $0.4 \le \mu \le 0.8$, $0.001 \le D \le 0.005$, $0 \le \kappa \le 0.002$, $0 \le \lambda \le 0.2$ and $0 \le \eta \le 0.04$. When the two plasmids are incompatible, their relative abundances in each species, referred to as $x$, $y$, exhibit a strong negative correlation and are constrained by $x + y < 1$, suggesting that these two plasmids cannot coexist stably in the same host cell.

(**D**) In testing the persistence potential, the species were distributed into multiple niches. Each niche had its own carrying capacity, and different species within the same niche competed with each other.

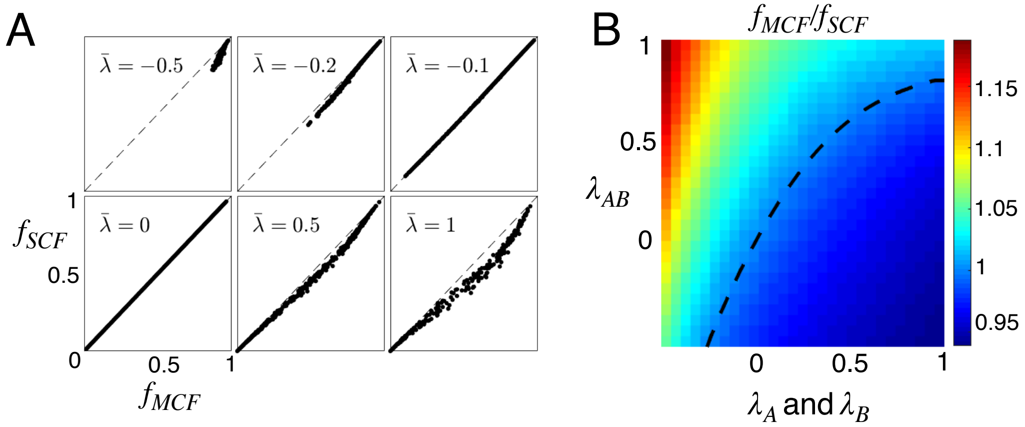


**Fig. S2. Systematic discrepancy between MCF and SCF.**

(**A**) For single-MCF communities, the discrepancy increases with higher burden or benefit. One thousand numerical simulations were performed in each of the six tests, with all the parameters in MCF and SCF randomized in the range of $1 \le m \le 50$, $1.6 \times 10^{-3} \le \kappa \le 2.4 \times 10^{-3}$, $1.6 \times 10^{-2} \le D \le 2.4 \times 10^{-2}$, $0 \le \eta \le 0.02$, and $0.3 \le \mu \le 0.8$. The steady-state abundance of the mobile element, defined as the fraction of MGE-carrying cells in the total population, was calculated as $f_{MCF}$ or $f_{SCF}$.

(**B**) For two-MGE communities, the discrepancy depends on their individual fitness costs ($\lambda_A$ and $\lambda_B$) as well as their combined effect ($\lambda_{AB}$). The parameters used in the simulations are $\kappa = 0.001$, $D = 0.005$, $\mu = 0.3$, $\eta = 0.01$. The ratio of $f_{MCF}$ to $f_{SCF}$ was calculated as the output. The conditions under which MCF and SCF generate the same MGE abundance ($\frac{f_{MCF}}{f_{SCF}} = 1$) are shown in black dashed curve.

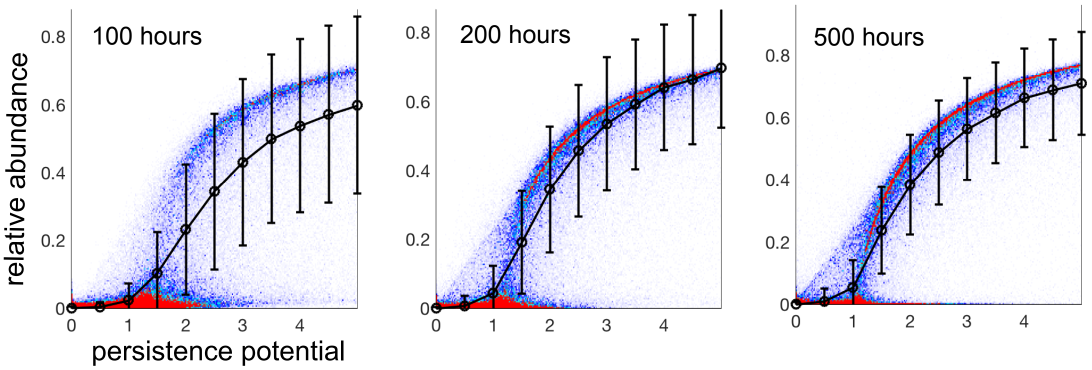


**Fig. S3. $\omega$ predicts temporal abundance of MGEs before the communities reach steady states.** 50,000 simulations were performed with 1-10 populations, 1-10 MGEs, and randomized parameters in the range of $0.1 \leq \mu \leq 0.8, 0.01 \leq D \leq 0.05$, $0 \leq \kappa \leq 0.1$, $0 \leq \lambda \leq 0.2$. For each simulation, the communities were assembled into a random number of niches. Within each niche, populations compete with each other. Each simulation was initialized with random abundances of populations and MGEs. The persistence potential $\omega$ of each MGE and its relative abundance in the entire community at different time points were then calculated. The $\omega$ range is divided into multiples bins with widths of 0.5. The standard deviations of the relative abundance in each bin are shown as error bars.

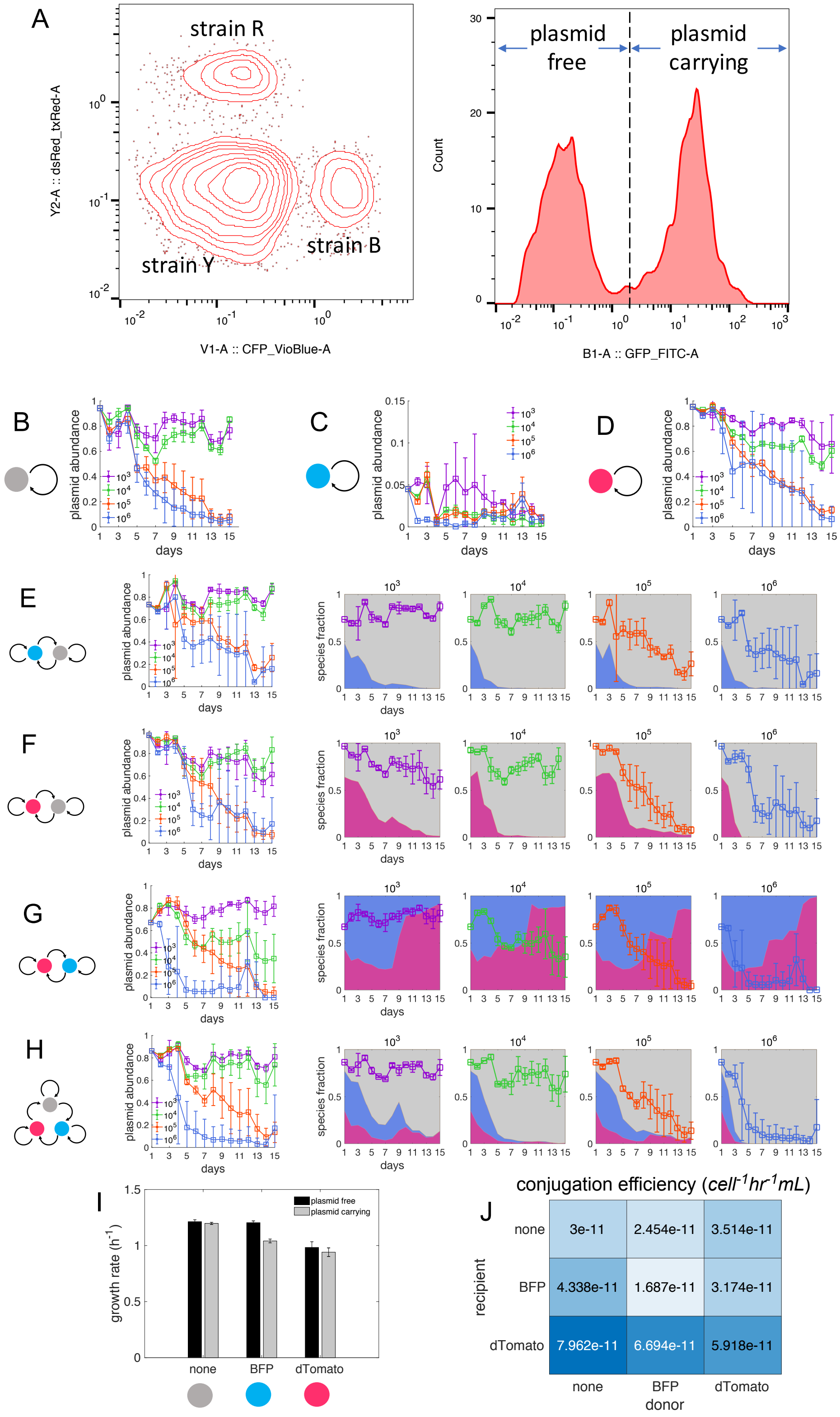

A
strain R
strain Y
strain B
Y2-A :: dsRed_txRed-A
V1-A :: CFP_VioBlue-A
plasmid free
plasmid carrying
Count
B1-A :: GFP_FITC-A
B
C
D
E
F
G
H
plasmid abundance
species fraction
days
I
growth rate (h-1)
plasmid free
plasmid carrying
none
BFP
dTomato
J
conjugation efficiency (cell-1hr-1mL)
recipient
donor
none
BFP
dTomato
3e-11
2.454e-11
3.514e-11
4.338e-11
1.687e-11
3.174e-11
7.962e-11
6.694e-11
5.918e-11

**Fig. S4. Long-term experiments and parameter estimations of the seven communities.**

(**A**) Determination of community composition and plasmid abundance using flow cytometry. V1 channel was used to detect BFP expressed by strain B, and Y2 channel was used to detect dTomato expressed by strain R. The cells with low V1 and Y2 signals represent strain Y. The plasmid-carrying cells, which express plasmid-borne GFP, were detected by channel B.

(**B-H**) The long-term dynamics of population composition and plasmid abundance in seven communities measured by flow cytometry. The filled circles stand for three *E. coli* strains. The strain expressing BFP chromosomally is shown in blue, the strain expressing dTomato is shown in red, and the non-fluorescent strain is shown in gray. The arrows shown in black represent the conjugation of the GFP-expressing plasmid K. Three communities were composed of single populations (B-D), three were composed of two members (E-G), and one was composed of all three members (H). Four daily dilution ratios ($10^3$, $10^4$, $10^5$m and $10^6$) were applied to each community. The plasmid dynamics are shown in line plots. The error bars stand for the standard deviations of three replicates. The composition dynamics are shown in colored areas.

(**I**) The growth rates of plasmid-free and plasmid-carrying cells of the three strains. The growth rates were calculated as the rate constants of the exponential phase. The error bars stand for the standard deviations of six replicates.

(**J**) The conjugation efficiencies among the three strains. Nine donor-recipient pairs were obtained, and the values of their conjugation rates are shown in the color map.

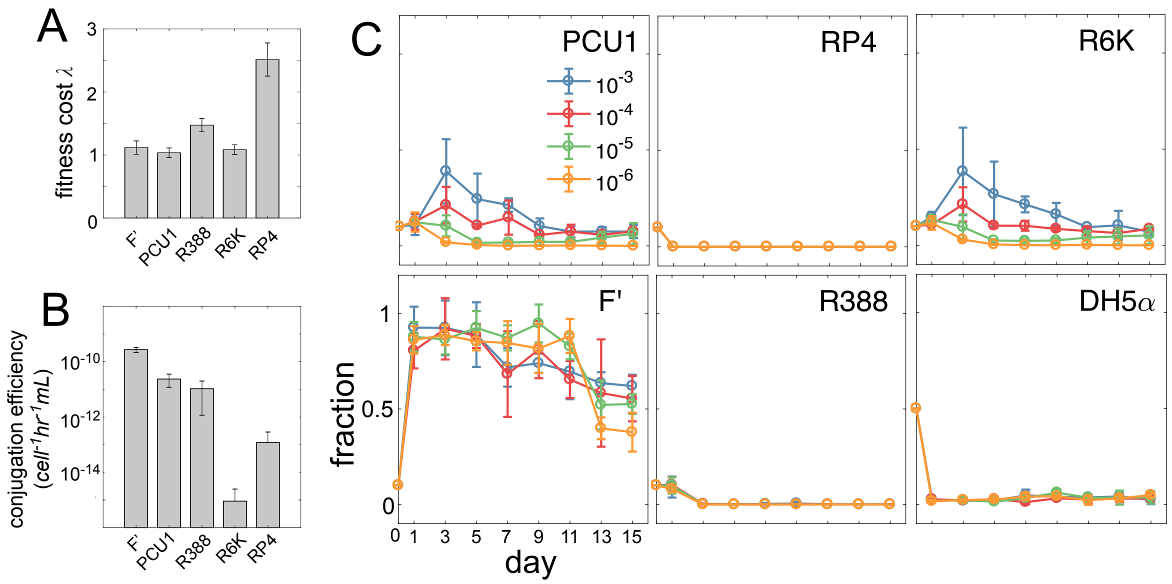


**Fig. S5. Long-term experiments and parameter estimations of the five-plasmid communities.**

(**A**) The fitness costs of the five conjugative plasmids in the strain MG1655. The error bars stand for the standard deviations of six replicates.

(**B**) The conjugation efficiencies of the five conjugative plasmids between MG1655 cells. The error bars stand for the standard deviations of three replicates.

(**C**) The long-term dynamics of population composition and plasmid abundance in the five-plasmid communities measured by plating. Four daily dilution ratios ($10^3$, $10^4$, $10^5$, and $10^6$) were applied to each community. The relative abundances of the plasmids (F', PCU1, R388, R6K, and RP4) and the strain DH5α are shown in line plots. The error bars stand for the standard deviations of three replicates.

#### References


1 Frost, L. S., Leplae, R., Summers, A. O. & Toussaint, A. Mobile genetic elements: the agents of open source evolution. *Nature Reviews Microbiology* **3**, 722 (2005).

2 Ochman, H., Lawrence, J. & Groisman, E. Lateral gene transfer and the nature of bacterial innovation. *Nature* **405**, 299-304 (2000).

3 Jørgensen, T. S., Xu, Z., Hansen, M. A., Sørensen, S. J. & Hansen, L. H. Hundreds of circular novel plasmids and DNA elements identified in a rat cecum metamobilome. *PloS one* **9**, e87924 (2014).

4 Kav, A. B. *et al.* Insights into the bovine rumen plasmidome. *Proceedings of the National Academy of Sciences* **109**, 5452-5457 (2012).

5 Zhang, T., Zhang, X.-X. & Ye, L. Plasmid metagenome reveals high levels of antibiotic resistance genes and mobile genetic elements in activated sludge. *PloS one* **6**, e26041 (2011).

6 Dahlberg, C., Linberg, C., Torsvik, V. L. & Hermansson, M. Conjugative plasmids isolated from bacteria in marine environments show various degrees of homology to each other and are not closely related to well-characterized plasmids. *Appl. Environ. Microbiol.* **63**, 4692-4697 (1997).

7 Brito, I. L. *et al.* Mobile genes in the human microbiome are structured from global to individual scales. *Nature* **535**, 435 (2016).

8 Dennis, J. J. The evolution of IncP catabolic plasmids. *Current Opinion in Biotechnology* **16**, 291-298 (2005).

9 Pilla, G. & Tang, C. M. Going around in circles: virulence plasmids in enteric pathogens. *Nature Reviews Microbiology* **16**, 484-495 (2018).

10 Mercer, R. *et al.* Genetic determinants of heat resistance in Escherichia coli. *Frontiers in microbiology* **6**, 932 (2015).

11 Broaders, E., O'Brien, C., Gahan, C. G. & Marchesi, J. R. Evidence for plasmid-mediated salt tolerance in the human gut microbiome and potential mechanisms. *FEMS microbiology ecology* **92** (2016).

12 Stokes, H. W. & Gillings, M. R. Gene flow, mobile genetic elements and the recruitment of antibiotic resistance genes into Gram-negative pathogens. *FEMS microbiology reviews* **35**, 790-819 (2011).

13 Ferreiro, A., Crook, N., Gasparrini, A. J. & Dantas, G. Multiscale evolutionary dynamics of host-associated microbiomes. *Cell* **172**, 1216-1227 (2018).

14 Iwasaki, W. & Takagi, T. Rapid pathway evolution facilitated by horizontal gene transfers across prokaryotic lineages. *PLoS genetics* **5**, e1000402 (2009).

15 Heuer, H. & Smalla, K. Plasmids foster diversification and adaptation of bacterial populations in soil. *FEMS microbiology reviews* **36**, 1083-1104 (2012).

16 Lopatkin, A. J. *et al.* Persistence and reversal of plasmid-mediated antibiotic resistance. *Nature communications* **8**, 1689 (2017).

17 Johnsen, P. J. *et al.* Factors affecting the reversal of antimicrobial-drug resistance. *The Lancet Infectious Diseases* **9**, 357-364 (2009).

18 Berg, P., Baltimore, D., Brenner, S., Roblin, R. O. & Singer, M. F. Asilomar conference on recombinant DNA molecules. *Science* **188**, 991-994 (1975).

19 Wright, O., Delmans, M., Stan, G.-B. & Ellis, T. GeneGuard: a modular plasmid system designed for biosafety. *ACS synthetic biology* **4**, 307-316 (2015).

20 Thiry, M. & Cingolani, D. Optimizing scale-up fermentation processes. *TRENDS in Biotechnology* **20**, 103-105 (2002).

21 Silva, F., Queiroz, J. A. & Domingues, F. C. Plasmid DNA fermentation strategies: influence on plasmid stability and cell physiology. *Applied microbiology and biotechnology* **93**, 2571-2580 (2012).

22 Slater, F. R., Bailey, M. J., Tett, A. J. & Turner, S. L. Progress towards understanding the fate of plasmids in bacterial communities. *FEMS Microbiology Ecology* **66**, 3-13 (2008).

23 Sørensen, S. J., Bailey, M., Hansen, L. H., Kroer, N. & Wuertz, S. Studying plasmid horizontal transfer in situ: a critical review. *Nature Reviews Microbiology* **3**, 700 (2005).

24 Bergstrom, C. T., Lipsitch, M. & Levin, B. R. Natural selection, infectious transfer and the existence conditions for bacterial plasmids. *Genetics* **155**, 1505-1519 (2000).

25 Stewart, F. M. & Levin, B. R. The population biology of bacterial plasmids: a priori conditions for the existence of conjugationally transmitted factors. *Genetics* **87**, 209-228 (1977).

26 Condit, R., Stewart, F. M. & Levin, B. R. The population biology of bacterial transposons: a priori conditions for maintenance as parasitic DNA. *The American Naturalist* **132**, 129-147 (1988).
27 Chao, L., Levin, B. R. & Stewart, F. M. A complex community in a simple habitat: an experimental study with bacteria and phage. *Ecology* **58**, 369-378 (1977).
28 Leclerc, Q. J., Lindsay, J. A. & Knight, G. M. Mathematical modelling to study the horizontal transfer of antimicrobial resistance genes in bacteria: current state of the field and recommendations. *Journal of the Royal Society Interface* **16**, 20190260 (2019).
29 Lozupone, C. A., Stombaugh, J. I., Gordon, J. I., Jansson, J. K. & Knight, R. Diversity, stability and resilience of the human gut microbiota. *Nature* **489**, 220 (2012).
30 Curtis, T. P., Sloan, W. T. & Scannell, J. W. Estimating prokaryotic diversity and its limits. *Proceedings of the National Academy of Sciences* **99**, 10494-10499 (2002).
31 Munck, C., Sheth, R. U., Freedberg, D. E. & Wang, H. H. Recording mobile DNA in the gut microbiota using an Escherichia coli CRISPR-Cas spacer acquisition platform. *Nature Communications* **11**, 1-11 (2020).
32 Martin-Serrano, J. & Neil, S. J. Host factors involved in retroviral budding and release. *Nature Reviews Microbiology* **9**, 519-531 (2011).
33 Lenski, R. E. & Levin, B. R. Constraints on the coevolution of bacteria and virulent phage: a model, some experiments, and predictions for natural communities. *The American Naturalist* **125**, 585-602 (1985).
34 Levin, B. R., Stewart, F. M. & Chao, L. Resource-limited growth, competition, and predation: a model and experimental studies with bacteria and bacteriophage. *The American Naturalist* **111**, 3-24 (1977).
35 Condit, R. The evolution of transposable elements: conditions for establishment in bacterial populations. *Evolution* **44**, 347-359 (1990).
36 Hall, J. P., Wood, A. J., Harrison, E. & Brockhurst, M. A. Source–sink plasmid transfer dynamics maintain gene mobility in soil bacterial communities. *Proceedings of the National Academy of Sciences* **113**, 8260-8265 (2016).
37 Loftie-Eaton, W. *et al.* Compensatory mutations improve general permissiveness to antibiotic resistance plasmids. *Nature ecology & evolution* **1**, 1354 (2017).
38 Hall, J. P., Williams, D., Paterson, S., Harrison, E. & Brockhurst, M. A. Positive selection inhibits gene mobilization and transfer in soil bacterial communities. *Nature ecology & evolution* **1**, 1348 (2017).
39 Harrison, E., Guymer, D., Spiers, A. J., Paterson, S. & Brockhurst, M. A. Parallel compensatory evolution stabilizes plasmids across the parasitism-mutualism continuum. *Current Biology* **25**, 2034-2039 (2015).
40 Kottara, A., Hall, J. P., Harrison, E. & Brockhurst, M. A. Variable plasmid fitness effects and mobile genetic element dynamics across Pseudomonas species. *FEMS microbiology ecology* **94**, fix172 (2017).
41 Dahlberg, C. & Chao, L. Amelioration of the cost of conjugative plasmid carriage in Eschericha coli K12. *Genetics* **165**, 1641-1649 (2003).
42 Fischer, E. A. *et al.* The IncI1 plasmid carrying the bla CTX-M-1 gene persists in in vitro culture of a Escherichia coli strain from broilers. *BMC microbiology* **14**, 77 (2014).
43 Porse, A., Schønning, K., Munck, C. & Sommer, M. O. Survival and evolution of a large multidrug resistance plasmid in new clinical bacterial hosts. *Molecular biology and evolution* **33**, 2860-2873 (2016).

44 Bohannan, B. J. & Lenski, R. E. Effect of resource enrichment on a chemostat community of bacteria and bacteriophage. *Ecology* **78**, 2303-2315 (1997).
45 Jones, G. W., Baines, L. & Genthner, F. J. Heterotrophic bacteria of the freshwater neuston and their ability to act as plasmid recipients under nutrient deprived conditions. *Microbial ecology* **22**, 15-25 (1991).
46 Winter, S. *et al.* Host-Derived Nitrate Boosts Growth of E. coli in the Inflamed Gut. *Science* **339**, 708-711 (2013).
47 Stecher, B. *et al.* Gut inflammation can boost horizontal gene transfer between pathogenic and commensal Enterobacteriaceae. *Proceedings of the National Academy of Sciences* **109**, 1269-1274 (2012).
48 Wotzka, S. Y., Nguyen, B. D. & Hardt, W.-D. Salmonella Typhimurium diarrhea reveals basic principles of enteropathogen infection and disease-promoted DNA exchange. *Cell host & microbe* **21**, 443-454 (2017).
49 Lopatkin, A. J. *et al.* Antibiotics as a selective driver for conjugation dynamics. *Nature microbiology* **1**, 16044 (2016).
50 Korem, T. *et al.* Growth dynamics of gut microbiota in health and disease inferred from single metagenomic samples. *Science* **349**, 1101-1106 (2015).
51 Arnoldini, M., Cremer, J. & Hwa, T. Bacterial growth, flow, and mixing shape human gut microbiota density and composition. *Gut microbes* **9**, 559-566 (2018).
52 Dionisio, F., Matic, I., Radman, M., Rodrigues, O. R. & Taddei, F. Plasmids spread very fast in heterogeneous bacterial communities. *Genetics* **162**, 1525-1532 (2002).
53 Mitreva, M. & Consortium, H. M. P. Structure, function and diversity of the healthy human microbiome. *Nature* **486**, 207-214 (2012).
54 David, L. A. *et al.* Diet rapidly and reproducibly alters the human gut microbiome. *Nature* **505**, 559 (2014).
55 Yatsunenko, T. *et al.* Human gut microbiome viewed across age and geography. *Nature* **486**, 222-227 (2012).
56 Willing, B. P., Russell, S. L. & Finlay, B. B. Shifting the balance: antibiotic effects on host–microbiota mutualism. *Nature Reviews Microbiology* **9**, 233-243 (2011).
57 Andersson, D. I. & Hughes, D. Antibiotic resistance and its cost: is it possible to reverse resistance? *Nature Reviews Microbiology* **8**, 260 (2010).
58 Sheth, R. U., Cabral, V., Chen, S. P. & Wang, H. H. Manipulating bacterial communities by in situ microbiome engineering. *Trends in Genetics* **32**, 189-200 (2016).
59 Ronda, C., Chen, S. P., Cabral, V., Yaung, S. J. & Wang, H. H. Metagenomic engineering of the mammalian gut microbiome in situ. *Nature methods* **16**, 167-170 (2019).
60 Gullberg, E. *et al.* Selection of resistant bacteria at very low antibiotic concentrations. *PLoS pathogens* **7** (2011).
61 Dimitriu, T. *et al.* Genetic information transfer promotes cooperation in bacteria. *Proceedings of the National Academy of Sciences* **111**, 11103-11108 (2014).
62 Balagaddé, F. K., You, L., Hansen, C. L., Arnold, F. H. & Quake, S. R. Long-term monitoring of bacteria undergoing programmed population control in a microchemostat. *Science* **309**, 137-140 (2005).
